# Dissecting neuron-specific functions of circadian genes using modified cell-specific CRISPR approaches

**DOI:** 10.1101/2023.02.23.529279

**Authors:** Shlesha Richhariya, Daniel Shin, Jasmine Quynh Le, Michael Rosbash

## Abstract

Circadian behavioral rhythms in *Drosophila melanogaster* are regulated by about 75 pairs of brain neurons. They all express the core clock genes but have distinct functions and gene expression profiles. To understand the importance of these distinct molecular programs, neuron-specific gene manipulations are essential. Although RNAi based methods are standard to manipulate gene expression in a cell-specific manner, they are often ineffective, especially in assays involving smaller numbers of neurons or weaker Gal4 drivers. We and others recently exploited a neuron-specific CRISPR-based method to mutagenize genes within circadian neurons. Here we further explore this approach to mutagenize three well-studied clock genes: the transcription factor gene *vrille,* the photoreceptor gene *Cryptochrome* (*cry*) and the neuropeptide gene *Pdf*. The CRISPR-based strategy not only reproduced their known phenotypes but also assigned *cry* function for different light mediated phenotypes to discrete, different subsets of clock neurons. We further tested two recently published methods for temporal regulation in adult neurons, inducible Cas9 and auxin-inducible gene expression system (AGES). The results were not identical, but both approaches successfully showed that the adult-specific knockout of the neuropeptide *Pdf* reproduces the canonical loss-of-function mutant phenotypes. In summary, a CRISPR-based strategy is a highly effective, reliable, and general method to temporally manipulate gene function in specific adult neurons.

**Significance statement:** Most animals have specific brain neurons that regulate sleep-wake cycles and other aspects of circadian behavior. *Drosophila* has only about 150 of these clock neurons. Despite their small numbers, they have remarkably diverse anatomy and gene expression profiles. To address the different functions of these neurons, we used highly specific and efficient CRISPR-based methods to create cell type-specific disruptions of three traditional circadian genes. We were able to assign the function of the photoreceptor cryptochrome to two tiny subsets of clock neurons. In addition, two independent methods assigned the neuropeptide PDF to the adult stage. In summary, we find that the CRISPR-based methods are very efficient at studying adult specific functions of genes in small, discrete sets of neurons.

## Introduction

Organismal behavior involves sensing the environment, internal processing, and motor actions. In multicellular organisms, this process is regulated largely by cells of the nervous system which often have defined roles as parts of bigger networks and circuits. One behavior that is almost ubiquitous to organisms on earth is aligning activity bouts to the daily rotations of the earth. The underlying mechanism is circadian rhythmicity.

In *Drosophila*, circadian clocks tick away in about 75 pairs of central brain clock neurons. These neurons all express the core circadian machinery, which includes the transcription factors CLOCK/CYCLE and the circadian inhibitory proteins PER/TIM. Rhythmic expression of these genes is important for maintaining circadian rhythmicity, and together they form a core transcription-translation feedback loop whose activity can be entrained by environmental cues such as light (1–3). Clock neurons also express several other proteins, such as the transcription factors VRILLE (VRI) and PDP1; their rhythmic expression is important for circadian behavior (4, 5).

Many additional genes are specifically expressed within clock neurons and also contribute to rhythmic behavior. Pigment dispersing factor (*Pdf*) encodes a circadian neuropeptide expressed exclusively within the small and large LNvs; *Pdf* mutants are arrhythmic and show increased sleep (6, 7). Another gene expressed by roughly half the clock neurons coding for the photoreceptor cryptochrome is *cry* (8–11). Wild-type flies are arrhythmic in constant light, whereas *cry* mutants are rhythmic in this condition (12). Despite the specificity in expression and probably function, few studies have explored the cell type-specific functions of these genes (13), and the cellular focus of many of circadian genes remains unknown.

Functional roles of specific circadian neurons have been broadly associated with the 7-8 subgroups based on anatomical location. For example, the PDF positive small LNvs are key regulators of rhythmicity under constant darkness and regulate morning activity. Another group of neurons, the 6 LNds along with the 5^th^ sLNv, regulate evening activity and thus were labelled evening cells (14–17). Yet other clock neuron subgroups play accessory roles in regulating rhythmicity and even sleep. They include the PDF-positive large LNvs, the DN1s, the DN2s, the LPNs and the mysterious DN3s (18–22)

Recent single cell sequencing indicates at least 17 distinct subgroups, about twice as many as originally identified based on anatomical location (23). It is likely that many of these new subgroups are also functionally distinct. Fortunately, many of them can now be accessed through judicious use of the Gal4/UAS system (24, 25), which allows for cell type-specific perturbations. This is because there are multiple Gal4 and split Gal4 drivers available to access specific circadian neurons (26, 27).

These drivers can be combined with the standard method of UAS-RNA interference (RNAi) to knockdown gene expression. Although this approach works well in many cases and has been invaluable for cell type-specific perturbations, it is often variable and usually induces partial knockdown (28, 29). Fortunately, it is now possible to disrupt genes completely and in the same cell-specific manner using CRISPR-Cas9-based gene editing (30, 31). Cas9 is an enzyme that induces a double stranded break at a specific genomic location determined by the guideRNA (gRNA) sequence (32). The error prone repair machinery endogenous to all cells usually leads to mutations known as indels (insertions-deletions). These range from a missense mutation to a small addition or deletion causing frameshift mutations, often disrupting protein function (31).

Earlier adaptations of this method achieved cell specificity by expressing the Cas9 protein in a Gal4 dependent manner while ubiquitously expressing a single guide RNA against a gene of interest. The method was found to be more consistent and effective than RNAi in inducing loss of function phenotypes (29, 33). More recent developments such as multiplexing guides against a gene in an array under UAS control further improved both the efficiency and cell type-specificity (33, 34). This improved method worked well for disrupting the functions of the core clock genes *period* (*per*) and *timeless* (*tim*) in circadian neurons (35, 36) and it was more recently extended to the disruption of genes encoding G-protein coupled receptors in these neurons (37).

Here, we implement this method to further understand the roles of three circadian behavior-associated genes: *vri, cry* and *Pdf*. We chose to generate a CRISPR-based line against *vri* because there are no successful RNAi data in the literature, and we chose *cry* and *Pdf* to address precise cell type-specific functions. Lastly, the CRISPR/Cas9 strategy was combined with two recent methods for temporal regulation, which successfully restricted gene editing and the resulting phenotype to specific neurons in just the adult stage. The results taken together substantially add to previous circadian analyses and identify specific subsets of clock neurons within which these genes function to regulate behavior.

## Results

### CRISPR mediated loss of VRI from clock neurons leads to a short circadian period

Circadian behavioral rhythms are primarily regulated by a molecular program operating in clock neurons. One component of this program is the transcription factor VRI encoded by the essential gene *vrille (vri)*. Homozygous mutants are developmentally lethal (38), and hemizygous mutants are strongly rhythmic but with a shorter free-running circadian period (4). Cycling of VRI in clock neurons is likely important as constant over-expression in clock-expressing cells causes arrhythmicity and a longer circadian period (4). However, an RNAi against *vri* in clock neurons did not affect circadian behavior (39). We therefore developed a line with CRISPR/Cas9 based 3X-guide-RNAs targeting the *vri* gene under UAS-control (*UAS-vri-g*). When expressed along with the reduced expression *UAS-Cas9.P2* (29) in clock neurons and the *CLK856-Gal4* driver, there was potent cell type-specific loss of VRI-staining (Fig. 1A, B, S1A). Importantly, the effect was shown to be specific as only VRI levels in Gal4-labeled cells were impacted; VRI levels in the DN3 neurons were unaffected (Fig. S1A) as expected since most of them are not labeled by the *CLK856-Gal4* driver (23). These flies had normal behavior under 12h light: 12h dark (LD) conditions, but with a small evening peak advance compared to the control strains (Fig. S1B, C). Consistent with this phenotype, the flies were completely rhythmic in DD but with a significantly shorter circadian period (Fig. 1C-G). These phenotypes are identical to those of a recent *vri* mutant (40) indicating that they derive from the clock neurons.

**Fig. 1:**
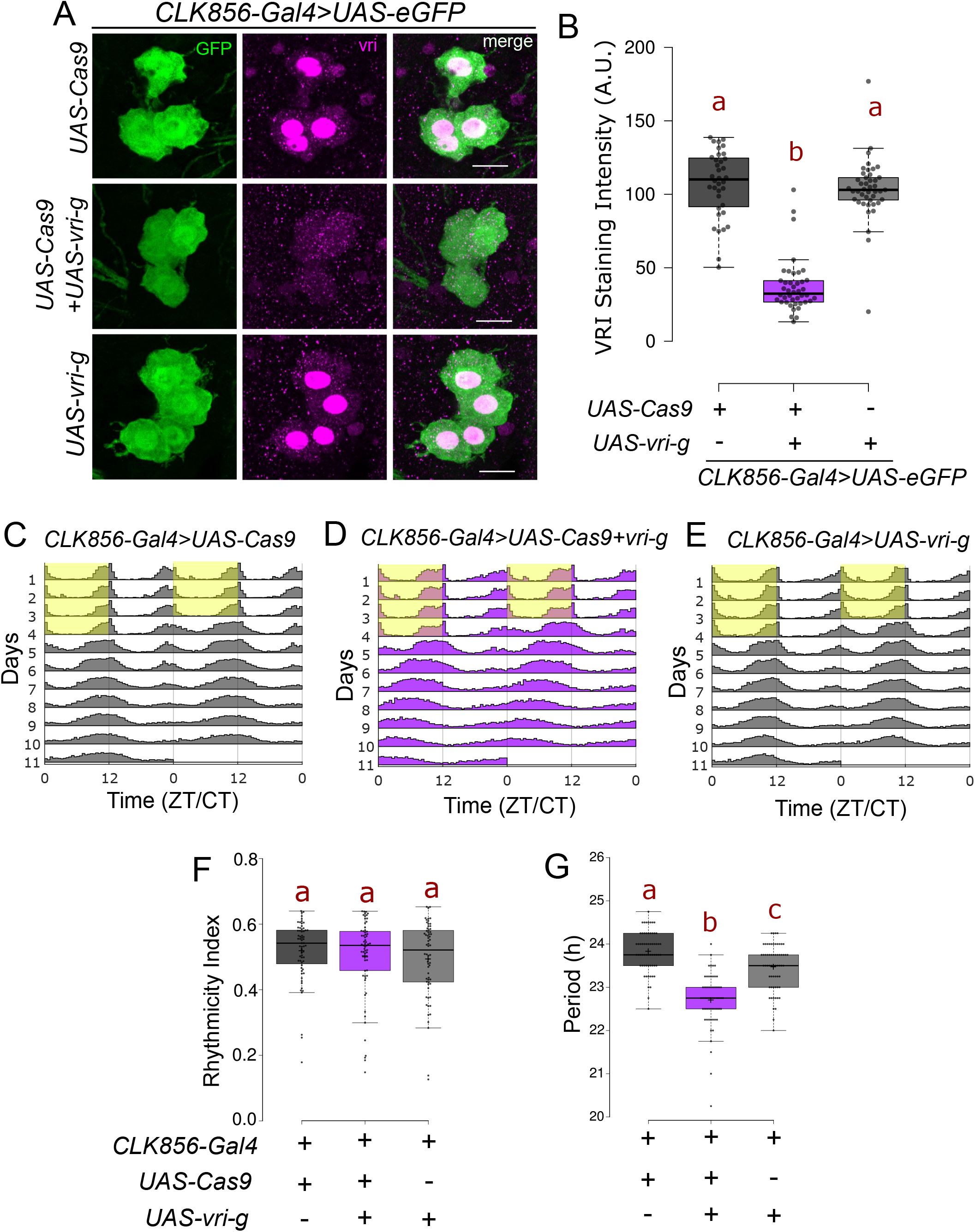
Loss of VRI from clock neurons leads to a shorter circadian period. **A**. Representative images of large LNV neurons stained for GFP and VRI at ZT19. There is loss of (nuclear) VRI staining in cells expressing both *UAS-Cas9* and UAS-3X-guides against *vri (UAS-vri-g)*. Some non-nuclear staining is visible in both control and mutant cells which could be due to background reactivity of the antibody. Scale bar represents 10µm. **B.** Quantification of VRI staining intensity from large LNv neurons represented as a boxplot, n ≥ 36 cells from at least 12 hemibrains per genotype, letters represent statistically distinct groups; p<0.001, Kruskal Wallis test followed by a post hoc Dunn’s test**. C-E**. Actograms represent double-plotted average activity of flies from an experiment across multiple days. Yellow panel indicates lights ON. **F.** Rhythmicity Index for individual flies plotted as a boxplot, n ≥ 62 per genotype from at least two independent experiments. No statistical difference was observed among the groups tested (p=0.6, Kruskal Wallis test). **G.** Free running period under constant darkness for individual rhythmic flies (RI>0.25) plotted as a boxplot, letters represent statistically distinct groups; p<0.01, Kruskal Wallis test followed by a post hoc Dunn’s test.

### CRY functions independently in two subsets of clock neurons

We next addressed the circadian gene *cry,* which encodes the photoreceptor Cryptochrome (CRY). Wild type flies are arrhythmic under constant bright light conditions, *cry* mutants in contrast remain rhythmic in constant light (12). This classic phenotype is reproduced when *cry* was mutated with *UAS-Cas9.P2* and 3X-gRNA against cry (*UAS-cry-g)* specifically in clock neurons using the *CLK856-Gal4* driver, which also led to loss of nearly all CRY staining from all clock neurons (Fig. S3). Whereas only about 10 percent of the control flies were rhythmic in constant light (Rhythmicity Index>0.25), 90 percent of the *cry-*mutated flies were rhythmic (Fig. 2A-C, G, Table S1). These flies were also rhythmic in constant darkness, like the controls (Fig. S2) and like traditional *cry* mutants (12).

**Fig. 2:**
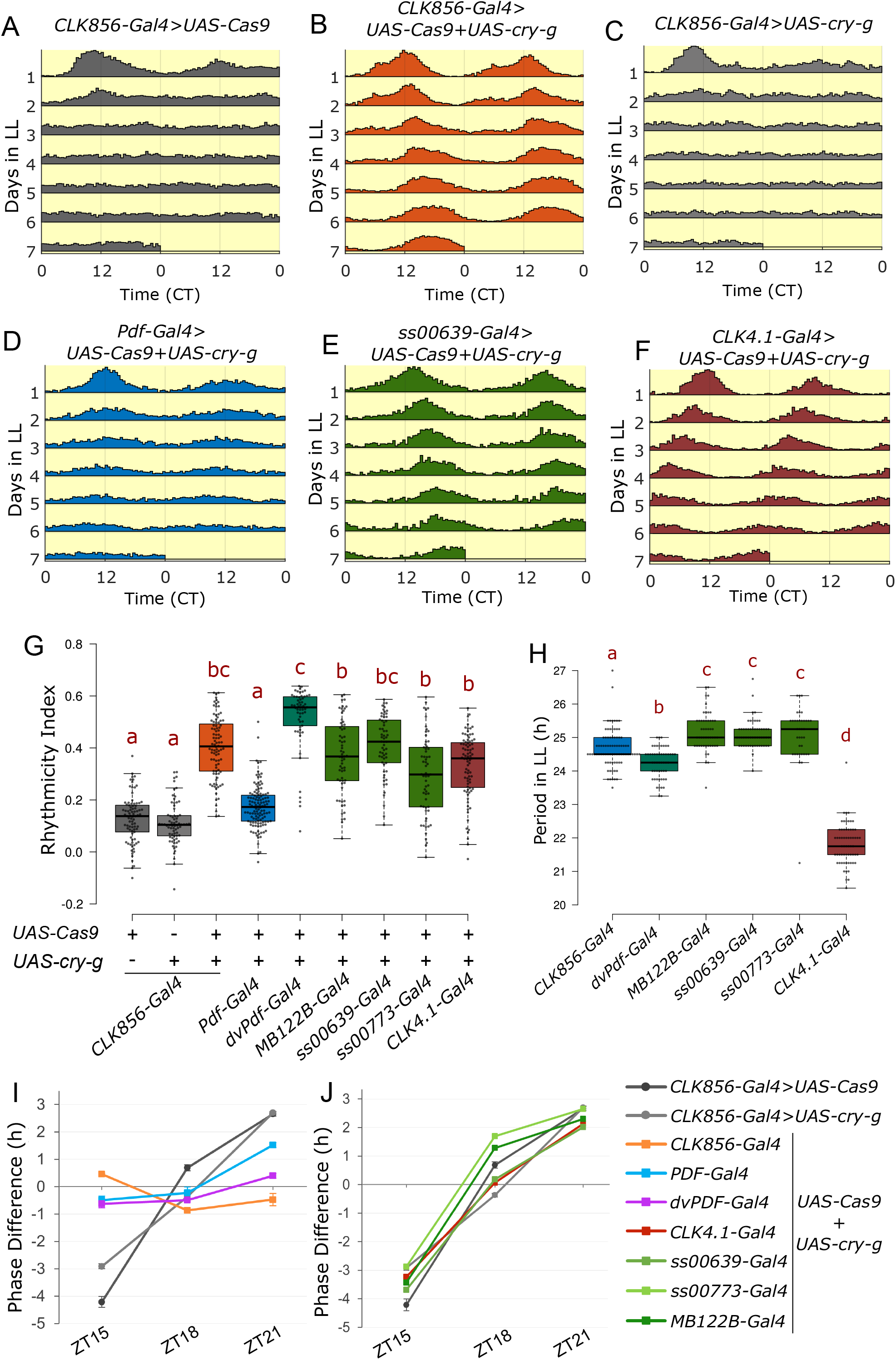
CRY function in two discrete subsets of clock neurons can regulate rhythmicity under constant light conditions. **A-F**. Actograms represent double-plotted average activity of flies from an experiment across multiple days in constant light. Flies expressing both *UAS-Cas9* and UAS-3X-guides against *cry (UAS-cry-g)* in clock neurons are rhythmic in constant light conditions. **G**. Rhythmicity Index of individual flies quantified for LL2-7 represented by a boxplot, n ≥ 51 per genotype from at least two independent experiments. Letters represent statistically distinct groups; p<0.001, Kruskal Wallis test followed by a post hoc Dunn’s test. **H**. Period under constant light for individual rhythmic flies (Rhythmicity Index > 0.25) quantified for LL2-7 represented by a boxplot, letters represent statistically distinct groups; p<0.01, Kruskal Wallis test followed by a post hoc Dunn’s test. **I-J.** Phase shifts in hours in response to a 60min light pulse at indicated times (n ≥ 26). Phase difference plotted as means ± SEM of differences from DD3-5. Same control flies are plotted in I and J.

As *CLK856-Gal4* is expressed in most clock neurons, we then asked if the constant light phenotype could be assigned to a more specific subset of circadian neurons. CRISPR-mediated cell type-specific mutagenesis of *cry* led to loss of CRY staining from specific neurons (Fig. S3). All PDF neurons are CRY-positive (8, 11), but disrupting *cry* in PDF cells with the *Pdf-Gal4* (7) had only a very small effect on constant light arrhythmicity (Fig. 2D, G, Table S1). In contrast, flies in which *cry* was mutated in the cells marked by the *dvPdf-Gal4* were fully rhythmic, indistinguishable from CRY loss in all circadian neurons (Fig. 2G, Table S1). This is interesting because the *dvPdf-Gal4* labels only 5 more cells per hemibrain than *Pdf-Gal4*, and only 2 of these 5 cells are CRY-positive: the 5^th^ sLNv and the one ITP-positive LNd (41, 42).

Ongoing work in our lab has characterized two Janelia split-Gal4 drivers, *ss00639-GAL4* and *ss00773-Gal4*. They both specifically label one CRY^+^ LNd and the 5^th^ sLNv in addition to very few additional cells (Fig. S4). Expression of CRISPR reagents against *cry* with the *ss00639-Gal4* driver also caused complete rhythmicity in constant light; more than 90% of the flies were rhythmic (Fig. 2E, G, Table S1). Similar but less strong results were seen with *ss00773-Gal4* (Fig 2G, Table S1). A third Gal4 driver, *MB122B-Gal4*, also expresses in these same two cells along with two additional LNd cells (43, 44), and mutating *cry* in cells labelled by this driver resulted in about 80% rhythmic flies in constant light (Fig. 2G, Table S1).

In addition to the lateral cells, CRY is expressed in the DN1p neurons labeled by the *CLK4.1-Gal4* driver (19, 45). Mutating *cry* in cells labeled by this driver also resulted in potent rhythmicity; nearly 75% of these flies were rhythmic in constant light (Fig. 2F, G, Table S1). These data taken together identify two sets of circadian neurons that can regulate rhythmicity independently under constant light in the absence of CRY: two evening cells consisting of the 5^th^ sLNv and the single ITP-positive LNd as well as the DN1p neurons.

To address possible mechanisms underlying the different groups of clock neurons and their regulation of constant light rhythmicity, we examined the circadian period of all rhythmic genotypes. Interestingly*, cry* mutants in all clock neurons have a circadian period of 24.7±0.7h in constant light, about an hour longer than their constant darkness period of 23.6±0.08 h (Fig. 2H, S2E). Mutating CRY in the evening cells with any of the three Gal4s gave rise to a similar period length of greater than 24 h, ∼25h (Table S1, green boxes in Fig. 2H). Surprisingly, mutating CRY only in the DN1ps caused a dramatically shorter constant light period of ∼22h (Table S1, Fig. 2H). These data indicate that specific clock neurons mediate the constant light effect and that there are interesting differences between the two discrete sets of relevant neurons (See Discussion).

In addition to being rhythmic under constant light, *cry* mutants are also deficient in their response to light pulses (46, 47). Control flies experience a delay phase shift in response to a light pulse in the early night at ZT15, an advance phase in response to a light pulse in the late night at ZT21 but little change with a light pulse in the middle of the night at ZT18 (46). The unresponsive *cry* mutant phenotype was reproduced by loss of CRY function specifically in clock neurons (Fig. 2I). Interestingly, flies with loss of CRY from PDF neurons were also completely deficient in the ZT15 light pulse-mediated phase delay but still maintained some phase advance induced by the ZT21 light pulse. In addition, when CRY function was perturbed in both PDF neurons and evening cells using the dvPDF-Gal4, the phase advance as well as the phase delay response was abolished (Fig. 2I). However, loss of CRY either in the evening cells alone or in the DN1p neurons did not affect the phase response (Fig. 2J). These data further indicate that CRY functions in discrete sets of clock neurons to mediate diverse light responses (see Discussion).

### PDF regulates activity and sleep from both the large and small LNvs

A key circadian molecule beyond the core clock is the neuropeptide pigment dispersing factor (PDF); it is encoded by the gene *Pdf*. Null mutants of the gene (*Pdf^01^*) are largely arrhythmic in constant darkness (7). PDF is expressed in only 8 pairs of lateral neurons in the adult brain, four small (sLNvs) and four large (lLNvs) ventral lateral neurons per hemibrain. The lLNvs express very high levels of PDF, which can be easily visualized within their long arborizations. They extend to the optic lobes and to the other side of the brain. The sLNvs extend a long dorsal process, which is also marked by PDF staining (Fig. S5A, (48)).

A UAS-3X-gRNA line against *Pdf* (*UAS-Pdf-g*) in combination with the *Pdf-Gal4* driver that labels all PDF-producing neurons caused loss of PDF from the cell bodies and projections of most PDF neurons (Fig. 3A, B and S5B). There was however residual staining in some lLNvs (Fig 3A, bottom panel, S5B), which likely reflects some residual PDF due to its very high levels in these cells prior to the mutagenesis. Nonetheless, this strain was completely arrhythmic in constant darkness whereas the parental control strains had normal rhythms (Fig. S6A-C, F). The very few rhythmic flies also had a shorter period phenotype (Fig. S6I).

**Fig. 3:**
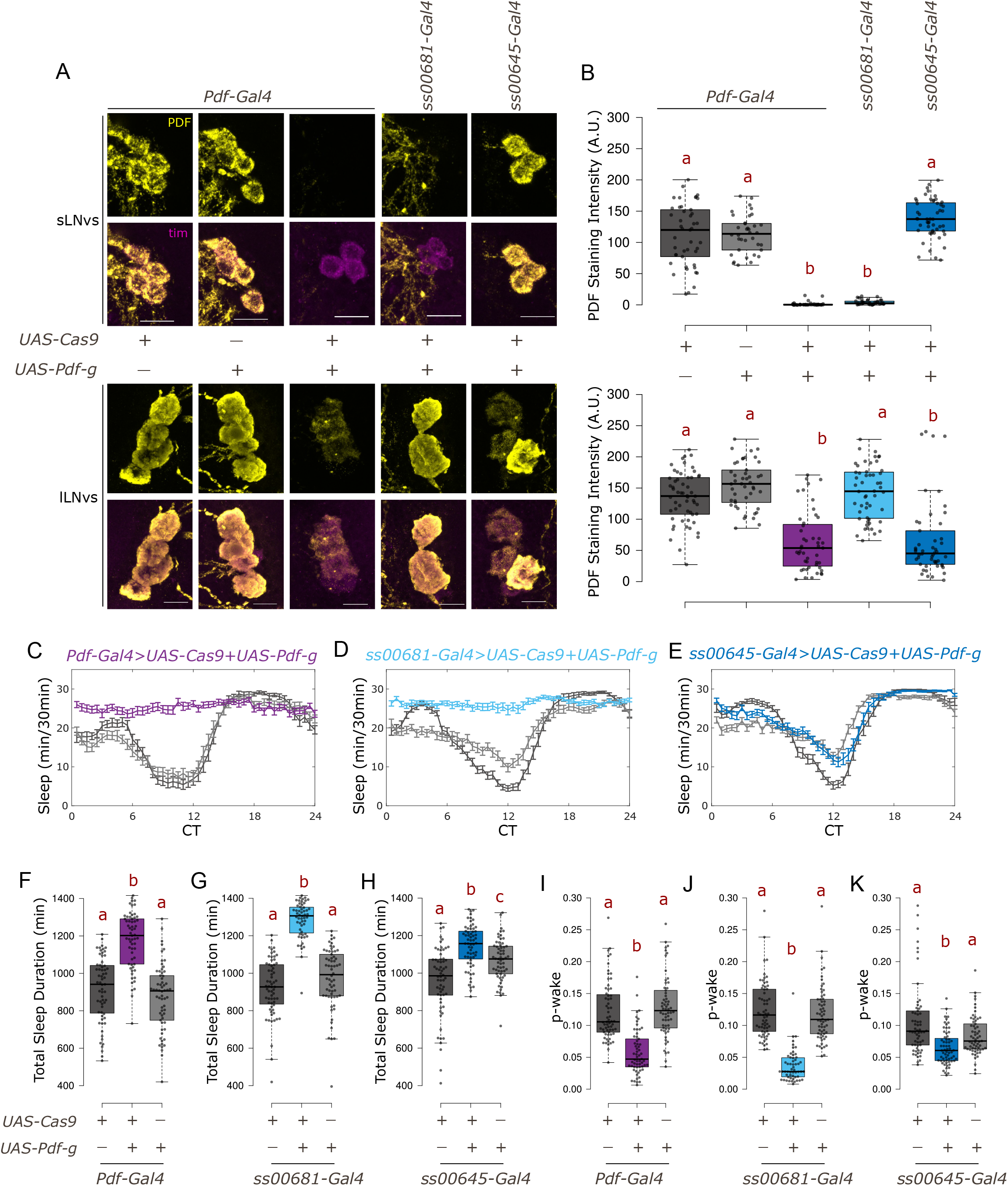
PDF from sLNvs maintains wakefulness under constant darkness. **A**. Representative images of sLNvs (top) and lLNvs (bottom) stained for TIM and PDF at ZT18. TIM labels all clock neurons. There is loss of PDF staining in cells expressing both *UAS-Cas9* and UAS-3X-guides against *Pdf (UAS-Pdf-g)*. Scale bar represents 10µm. **B**. Quantification of PDF staining intensity from sLNv (top) and lLNv (bottom) neurons represented with boxplots, n ≥ 36 cells from at least 11 hemibrains per genotype. Letters represent statistically distinct groups; p<0.001, Kruskal Wallis test followed by a post hoc Dunn’s test. **C-E**. Sleep plots representing average sleep of flies from DD1-4 in 30-minute bins. *UAS-Cas9* alone control is indicated in dark gray, *UAS-Pdf-g* alone control in light gray whereas the experimental genotype expressing both is represented by color indicated above the plots. **F-H**. Total sleep duration in a day for individual flies quantified (from DD1-4) represented by a boxplot, n ≥ 55 per genotype from at least two independent experiments. Letters represent statistically distinct groups, p<0.01; Kruskal Wallis test followed by a post hoc Dunn’s test. **I-K**. P-wake over a 24-hour period for individual flies quantified (from DD1-4) represented by a boxplot, letters represent statistically distinct groups; p<0.001, Kruskal Wallis test followed by a post hoc Dunn’s test.

We then used previously described split-Gal4 lines generated by the Rubin lab at Janelia (36) to specifically target only sLNvs or only lLNvs. The CRISPR-Cas9-based method was again highly effective in targeting PDF levels only in the cells targeted by the respective Gal4s. Expression of the gRNA and Cas9 in sLNv-specific driver (*ss00681-Gal4*) led to near complete loss of PDF staining from 100% of the sLNvs examined and their dorsal processes, with no effect on the lLNv cell bodies or on their processes. Similarly, expressing the CRISPR reagents using the lLNv-specific driver (*ss00645-Gal4*) significantly reduced PDF expression from lLNv cell bodies and processes, without affecting PDF levels in the sLNvs or their dorsal process (Fig. 3A, B; S5B). However, the effect on the lLNvs was not as strong: about 20% of the neurons (1-2 neurons from about half the hemibrains) still had unchanged PDF staining. This is very likely because the *ss00645-Gal4* variably labels 3-4 lLNvs per hemisphere. Notably however, there remains some residual staining in the lLNvs even with *Pdf-Gal4* (Fig. 3A, B-bottom panels), probably due to the higher levels of PDF in the lLNvs (see Discussion). As expected based on earlier studies (13), loss of PDF from the sLNvs but not from the lLNvs caused arrhythmicity in constant darkness (Fig. S6D-H).

Loss of PDF from both small and large PDF neurons also caused substantially increased sleep under constant darkness (Fig. 3C, F), and targeting the guides specifically to the small cells caused a similar sleep increase (Fig. 3D, G). This increase in sleep is not because of a lack of the ability to move as these flies are even more active than the controls when awake (Table S2). Although a role for PDF neurons in sleep and arousal has been described (6, 49), CRISPR-mediated loss of PDF specifically from the sLNvs here is the most direct evidence that the sLNvs regulate sleep. Loss of PDF from only the large cells also increased sleep but to a much smaller extent (Fig. 3E, H). Although the difference between the small and large cells may be due to the residual large cell PDF that escapes knockdown, or perhaps to the few lLNvs not labelled by the split-Gal4 driver, the very significant reduction in PDF signal from lLNv cell bodies and processes in most brains (Fig 3A-B, bottom panels; S5B) makes it more likely that the lLNvs have a smaller role in regulating sleep than the sLNvs in constant darkness.

An analysis of sleep structure indicated a substantial decrease in p-wake (the instantaneous probability that the flies will awaken from sleep), but no effect on p-doze (the instantaneous probability that awake flies will fall asleep (50)) (Fig. 3I-K, Table S2). These data are consistent with a wake-promoting role of PDF (6) and identify PDF from the sLNvs as the primary regulator.

### Temporally regulated CRISPR-Cas9 mutagenesis assigns PDF function to adult neurons

The function of several circadian genes has not yet been definitively assigned to the adult fly, so some of them might have a substantial developmental role. Traditional methods of temporal regulation of Gal4 activity such as GeneSwitch (51) or the temperature-sensitive Gal80 system (52) are not ideal for behavior experiments as they either involve toxic drugs or dramatic shifts in temperature, both of which severely impact the standard locomotor behavior assay (53, 54). Pairing with the temperature sensitive Gal80 system also resulted in leaky mutagenesis (33). We therefore assessed two more recently developed methods, one specific to CRISPR and the other more general, to mutagenize genes of interest adult-specifically.

The first was the inducible Cas9 method developed by Port et. al. (33). It is a UAS-based method: the Cas9 sequence is preceded by a GFP cassette, which is followed by a STOP codon flanked by FRT sites (hereafter called STOP-Cas9). Pairing with a heat-shock induced flp transgene allows for temporal control of mutagenesis as Cas9 expression occurs only post heat shock (Fig. S7A). We adapted this method for adult-specific induction by using three consecutive heat shocks (referred to below as “heat shock”; see Methods). Since these heat shocks are temporally separated by several days from the behavior analyses, the heat shocks are not expected to alter circadian behavior. The STOP-Cas9 transgene was combined with *Pdf-Gal4*, *UAS*-*Pdf-g,* a *UAS-mRFP* transgene to label the PDF cells as well as a heat shock induced flp encoding hsFLPG5 (Fig. S7A).

In control flies without *UAS*-*Pdf-g*, heat shock did not affect the levels of PDF (Fig. 4A, B, S7B) and did not negatively impact rhythmic behavior (Fig. 4C, D). Importantly, flies expressing all the transgenes including the UAS-gRNA but without heat shock showed no loss of PDF signal (Fig. 4A, B, S7B) and no defects in behavior (Fig. 4C-F); this indicates no leaky expression without heat shock. In the same flies two weeks post heat shock, PDF staining was lost from all sLNvs in ∼60% of the hemibrains analyzed, whereas some mosaicism was observed in the remaining 40% of the hemibrains; 1-2 neurons still expressed both GFP and PDF (Fig. 4A, B; S7B). A similar fraction of the heat shocked mutant flies, about 38 percent, were rhythmic in constant darkness (Fig. 4D, S7C), indicating an adult-specific requirement of PDF for rhythmicity. The ca. 60 percent arrhythmic flies also showed an adult-specific effect of PDF on sleep (Fig. 4E).

**Fig. 4:**
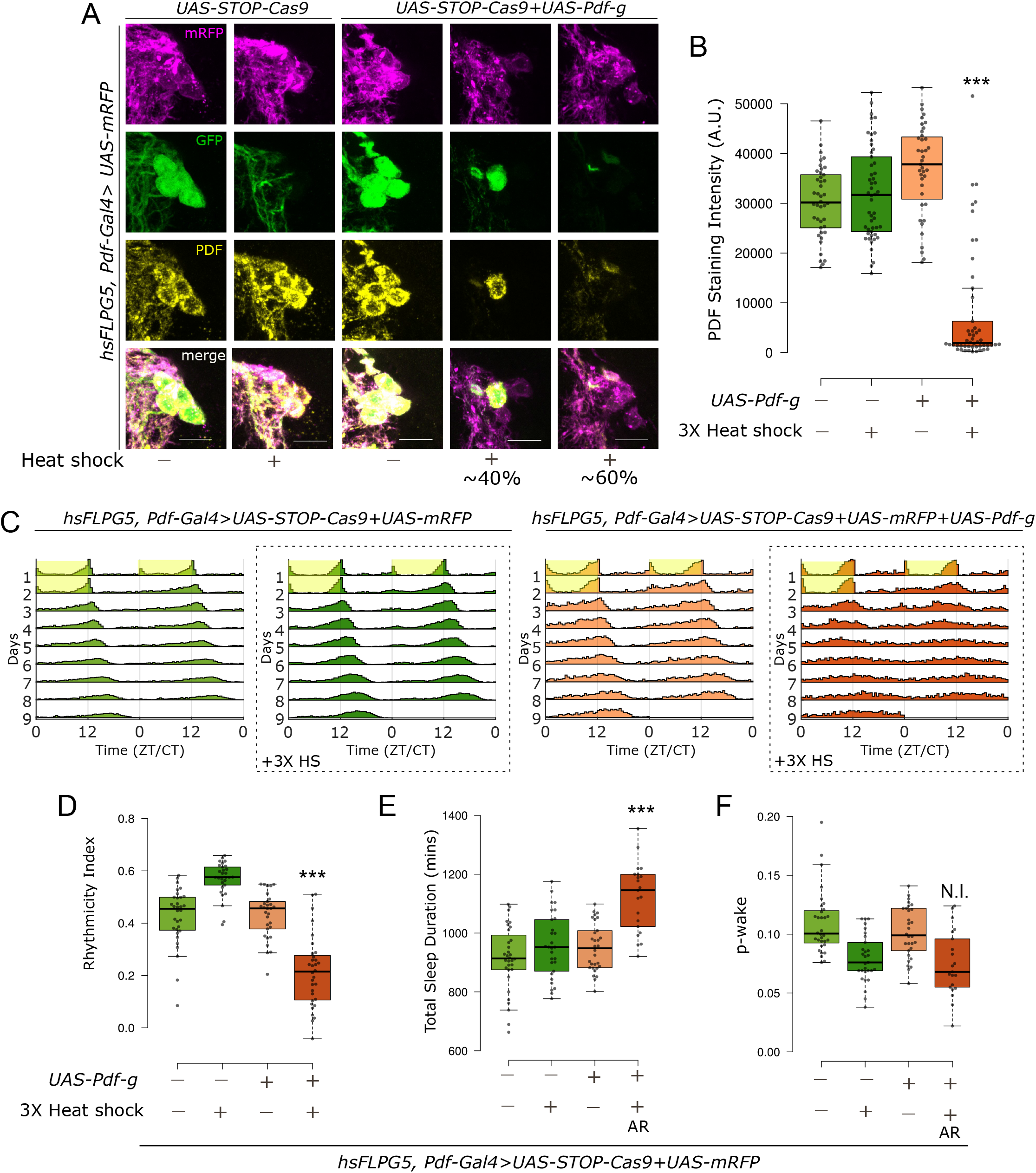
Adult-specific loss of PDF using an inducible Cas9 system results in loss of rhythmicity and increased sleep. **A**. Representative images of sLNvs stained for GFP, mRFP and PDF. 2 weeks post heat shock, there is loss of PDF staining in cells expressing both *UAS-Cas9* and *UAS*-*Pdf-g*. In about 60% of the hemibrains analyzed, PDF staining was lost from all sLNvs, whereas in the remaining 40% hemibrains, some mosaicism was observed as PDF staining was unchanged in 1-2 sLNvs. Scale bar represents 10µm. **B**. Quantification of PDF staining intensity from sLNvs represented with boxplots, n ≥ 39 cells from at least 12 hemibrains per genotype/condition. There is a significant effect of both variables on the staining intensity and a significant interaction when CRISPR reagents for inducible CRISPR for *Pdf* were combined with 3X heat shock (Two-way ANOVA; *** Interaction p<0.001). **C**. Actograms represent double-plotted average activity of flies from an experiment across multiple days. Yellow panel indicates lights ON. Panels in dotted boxes indicate heat-shocked flies. **D**. Rhythmicity Index for individual flies quantified (from DD2-7) represented by a boxplot, n ≥ 29 per genotype. There is a significant effect of both variables on rhythmicity and a significant interaction when CRISPR reagents for inducible CRISPR for *Pdf* were combined with 3X heat shock (Two-way ANOVA; *** Interaction p<0.001). **E**. Total sleep duration in a day for individual flies quantified for DD1-4 represented by a boxplot. For the group expressing both *UAS-Cas9* and *UAS-Pdf-g* with heat-shock, only data from the arrhythmic flies are shown and used for statistics (indicated by AR). Also see Fig. S5D. There is a significant effect of both variables on sleep duration and a significant interaction when CRISPR reagents for inducible CRISPR for *Pdf* were combined with 3X heat shock (Two-way ANOVA; *** Interaction p<0.001). **F**. p-wake over a 24-hour period for individual flies quantified (from DD1-4) represented by a boxplot. For the group expressing both *UAS-Cas9* and *UAS-Pdf-g* with heat-shock, only data from the arrhythmic flies are shown and used for statistics (indicated by AR). Also see Fig. S5D. Only heat-shock had a significant effect on p-wake and there was no interaction between the variables (Two-way ANOVA; N.I. = No Interaction, p=0.31).

The remaining rhythmic flies likely reflect mosaicism, as PDF from 1-2 neurons might be sufficient to maintain some level of rhythmicity. This is supported by the fact that the rhythmicity and sleep phenotypes were correlated in the mutant flies (Fig. S7D). Surprisingly, heat shock alone had a strong effect on the sleep structure and lowered p-wake significantly in the controls. Therefore, no conclusions could be drawn about the adult-specific effect of PDF in maintaining p-wake (Fig. 4F). These data indicate that the heat-shock based methods are compatible with studying circadian behavior but may be less suitable for studying sleep behavior.

We also combined CRISPR-Cas9 mutagenesis with a more general method of temporal regulation that affects the Gal4 and therefore controls all UAS-based strategies, the recently developed auxin-inducible gene expression system (AGES, (55)). It uses the traditional Gal4 repressor Gal80, which has been engineered to be regulated by auxin. Auxin is non-toxic and does not affect fly lifespan at the required concentrations (55). We combined the AGES system with the regular Cas9, *UAS-Cas9.P2,* along with *UAS-Pdf-g* and *UAS-mRFP* to label the LNvs with the *Pdf-Gal4* driver (Fig. S8A).

AGES sequesters GAL4 and thus inhibits expression from all UAS-transgenes without auxin feeding (Fig. S8A, (55)). Indeed, there is no to very low mRFP expression in cell bodies or processes without auxin feeding (Fig. 5A, S8B). However, expression must be non-zero, as some PDF mutagenesis is observed without auxin feeding (∼2-fold reduction in PDF levels, Fig. 5A, B). Not surprisingly, auxin feeding leads to a much bigger effect - near loss of all PDF from cell bodies (>10-fold reduction) as well as processes (Fig. 5A, B, S8B).

**Fig. 5:**
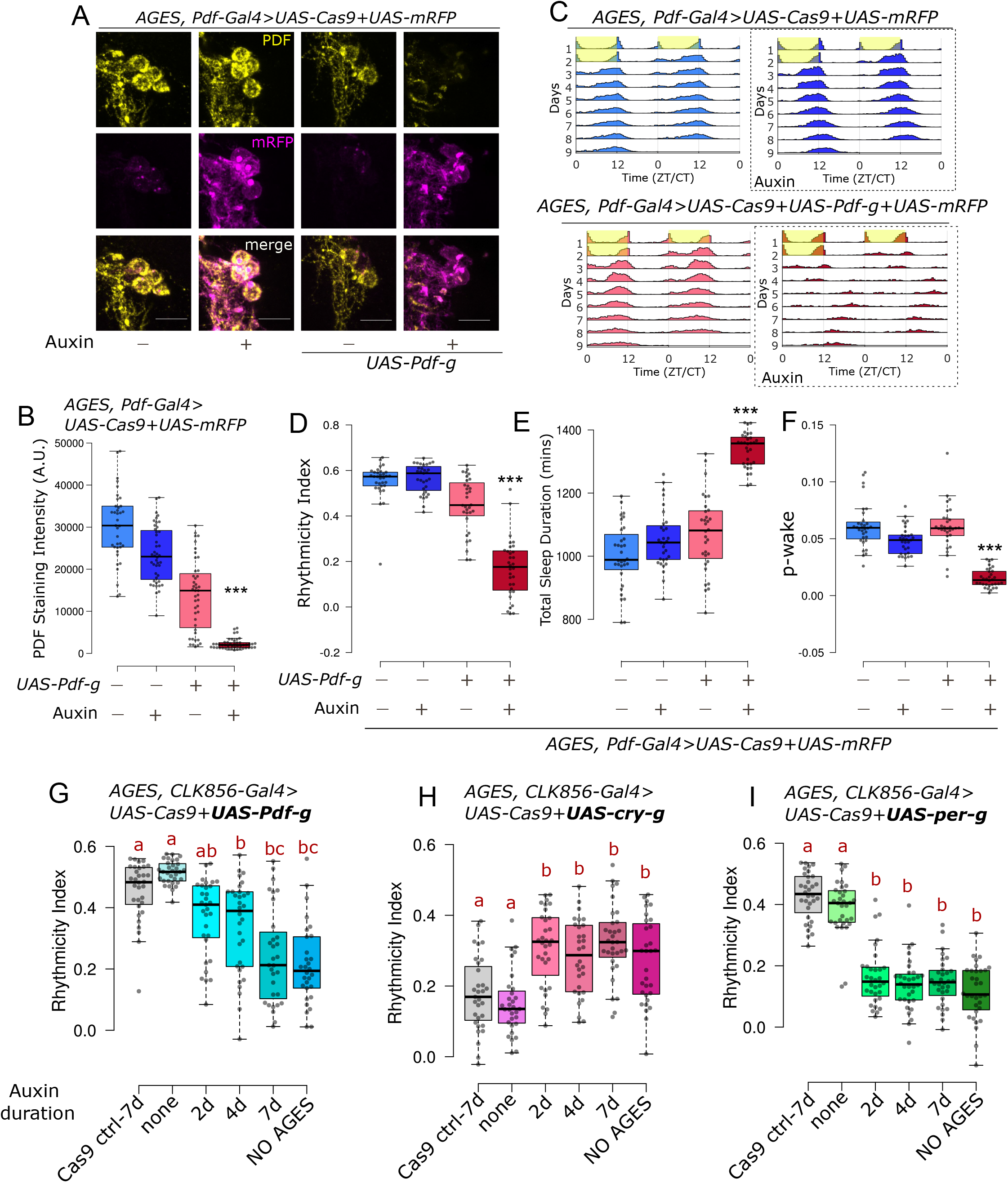
Adult-specific loss of PDF with AGES leads to loss of rhythmicity and wakefulness. **A**. Representative images of sLNvs stained for mRFP and PDF. Upon feeding 10mM auxin for 2 weeks, there is loss of PDF staining in cells expressing both *UAS-Cas9* and *UAS-Pdf-g*. Scale bar represents 10µm. **B**. Quantification of PDF staining intensity from sLNvs is represented with boxplots, n ≥ 39 cells from at least 11 hemibrains per genotype/condition. There is a significant effect of both variables on PDF staining intensity and a significant interaction when AGES and CRISPR reagents for *Pdf* were combined with auxin feeding (Two-way ANOVA; *** Interaction p<0.001). **C**. Actograms represent double-plotted average activity of flies from an experiment across multiple days. Yellow panels indicate lights ON. Panels in dotted boxes indicate auxin feeding. **D**. Rhythmicity Index for individual flies quantified (from DD2-7) represented by a boxplot, n ≥ 31 per genotype. There is a significant effect of both variables on rhythmicity and a significant interaction when AGES and CRISPR reagents for *Pdf* were combined with auxin feeding (Two-way ANOVA; *** Interaction p<0.001). **E**. Total sleep duration in a day for individual flies quantified (from DD1-4) represented by a boxplot. There is a significant effect of both variables on sleep duration and a significant interaction when AGES and CRISPR reagents for *Pdf* were combined with auxin feeding (Two-way ANOVA; *** Interaction p<0.001). **F**. p-wake over a 24-hour period for individual flies quantified (from DD1-4) represented by a boxplot. There is a significant effect of both variables on p-wake and a significant interaction when AGES and CRISPR reagents for *Pdf* were combined with auxin feeding (Two-way ANOVA; *** Interaction p<0.001). **G-I**. Rhythmicity Index for individual flies quantified (from DD2-6) represented by a boxplot, n ≥ 30 per genotype. Flies of the indicated genotypes were fed auxin for either 2-, 4- or 7-days and compared with flies that were fed no auxin, or with flies with mutagenesis throughout development (NO AGES). Letters represent statistically distinct groups;, Kruskal Wallis test followed by a post hoc Dunn’s test, p<0.003.

The small-scale leaky mutagenesis without auxin feeding had only a minor effect on rhythmicity and sleep (Fig. 5C, D, E). However, auxin feeding caused a very strong effect on both rhythmicity and sleep just like the effect on PDF staining (Fig. 5D, E, S8C), and adult-specific loss of PDF lowered p-wake (Fig. 5F). Auxin feeding alone in control flies had a very minor effect on PDF staining (20% reduction in cell bodies) and no effect on circadian behavior and sleep duration (Fig. 5A-E, S8B, C). There was also a significant interaction between auxin feeding and expression of CRISPR reagents against PDF on p-wake (p<0.001, Two-way ANOVA; Fig 5F). Combining AGES with the CRISPR system therefore indicate that most if not all aspects of PDF function on circadian behavior and sleep are adult-specific.

We then assayed the time required for adult-specific-CRISPR mediated mutagenesis. Flies expressing *UAS-Cas9.P2* and *UAS-Pdf-g* along with AGES were fed 10mM auxin for 2-, 4- and 7-days before testing for behavior on food without auxin. 2 days of auxin feeding did not significantly alter rhythmicity of the controls, but there was a gradual reduction in rhythmicity from 4 days to 7 days of auxin feeding (Fig. 5G). To further tease apart the required time for CRISPR mutagenesis from the effects of protein turnover, we used the same paradigm to test two proteins that undergo daily degradation – CRY and PER. For both proteins, 2 days of auxin feeding was sufficient to cause a phenotype indistinguishable from the perturbations through development (Fig. 5H, I), indicating that no more than 2 days are required for mutagenesis.

## Discussion

In this study we generated new UAS-3X-gRNA lines for neuron-specific perturbation of three key circadian genes and combined them with defined Gal4 drivers to assay their function in specific subsets of clock neurons. We could reproduce the mutant phenotypes of *vri, cry* and *Pdf* in known neurons and identified previously unknown cell type-specific functions of these genes. The approach worked very well with specific and weak Gal4 drivers, which allowed the discovery of two subsets of circadian neurons within which CRY acts independently to regulate rhythmicity under constant light conditions. Finally, these new reagents were compatible with existing tools and shown to be highly effective with two independent methods for temporal regulation. The results and lines described here will be of use to the circadian community, but we hope that the detailed validation of the method will appeal to a much wider population of fly neuroscientists and encourage them to switch from the standard but less dependable RNAi methodology to this more effective and consistent method for the spatial and temporal knockout of genes in adult fly neurons.

Our data show that this cell type-specific CRISPR method works well with highly expressed genes like *Pdf* and even with weak split-Gal4 drivers. It is also ideal for addressing the adult function of essential genes such as *vri*, by perturbing their expression levels and function in inessential cells as we show here for circadian neurons. In this context, the functions of many *Drosophila* genes are defined only in developmental contexts, even though these same genes are expressed again in adults. For example, we recently found that genes encoding GPCRs, transcription factors and cell surface molecules are broadly yet specifically expressed in clock and dopaminergic neurons. Studying their function in these specific adult neurons is beginning to help define the molecular underpinnings of neural circuits and behavior (23, 56). Towards these goals, we have generated and begun to use a library of guideRNA lines against all GPCRs expressed in *Drosophila* (37). Moreover, several lines with 2X-gRNA generated by Port and Boutros (33) are now publicly available. Such resources should also make this method a first choice for tissue and cell type-specific reverse genetic screens.

Another novel application of this method should be cell-specific gene-interaction studies. Combining multiple RNAi lines is not common, perhaps because of the suspected dilution effects of multiple UAS lines. In contrast, this CRISPR-Cas9 method does not require high expression levels, so combining gRNAs against multiple genes, for example to study enhancer and suppressor phenotypes, should be possible and would extend the approach in an interesting direction.

VRI operates as a part of the molecular transcription translation feedback loop that operates in clock neurons (57). Hemizygous mutants of *vri* are rhythmic with a short circadian period (4). Concurrently, a recent study generated a new homozygous mutant that has normal rhythms with a short circadian period despite dramatically reduced VRI levels in the clock neurons (40). These results are in complete agreement with ours (Fig. 1C-G). It is also possible that very low levels of VRI in clock neurons is sufficient to maintain rhythms like in the whole body mutant (40). Although unlikely in our view, this interpretation is consistent with an earlier study (58). It used cell-specific flip outs of a rescue transgene in a mutant *vri* background and found that VRI in PDF neurons regulates rhythmicity, but this result could have been influenced by the mutant background (58). Although over-expression of VRI also results in considerable loss of rhythms (4), simple loss of function via these CRISPR-Cas9-based methods is less prone to misinterpretation than gain-of-function experiments. In the future, guides can be targeted to specific subsets of circadian and non-circadian cells to further define the function of VRI in the regulation of development and behavior. The CRY photoreceptor along with the compound eyes and H-B eyelets are the three pathways through which flies entrain to environmental light cues (59). It is therefore not surprising that *cry* mutants are still able to light entrain normally. However, these mutants are rhythmic under constant bright light conditions, indicating a special role for CRY under prolonged light exposure conditions (12). CRY is expressed in several different clock neurons; they include the PDF neurons, a subset of evening cells (5^th^ sLNv+3 LNds) and a subset of the DN1ps (8). Interestingly, we find that CRY only functions in the evening cells and the DN1ps to regulate the response to constant light. This conclusion agrees with previous results indicating that evening cells (60, 61) and dorsal neurons (60, 62) are the principal drivers of rhythmicity under constant light conditions. QSM is another protein that mediates light input to the circadian clock, and flies mutant for QSM in dorsal neurons maintain rhythmicity in constant light (63), further emphasizing the role of dorsal neurons in mediating rhythmicity under constant light conditions.

Our results also agree with earlier results that PDF positive LNvs have no more than a minor role in responding to constant light (61, 62). Because the sLNvs are directly downstream of the light responsive H-B eyelets (64), these neurons may be constantly electrically stimulated and their molecular clock therefore poisoned by constant light even without CRY. In contrast, loss of CRY in either of the two other major subsets of CRY-positive neurons, the two evening cells (5^th^ sLNv, CRY^+^ and ITP^+^ LNd) as well as the DN1ps, is sufficient to maintain rhythmicity in constant light. These data indicate that CRY may be the primary source of light information in these neurons; perhaps they are critical for adapting to altered photoperiods such as longer summer days.

Interestingly, CRY functions in a different subset of clock neurons, the PDF neurons, to effect light pulse-mediated phase delays in the early night. Phase delays induced by a light pulse in the late night also requires CRY in PDF neurons but additionally requires CRY in evening neurons (Fig. 2I). A CRY requirement for phase responses in PDF neurons has been previously described (65), and a role for the evening neurons in phase response behavior has also been suggested (60). As neuronal activation of PDF neurons at different times of day can mimic a light-mediated phase response curve in a CRY independent manner (66), CRY photon capture may impact neuronal firing, a result consistent with direct observation (67), so altered electrical properties of the CRY mutants might also contribute to the observed light-mediated phenotypes.

Expression of *UAS-Cas9* and *UAS-Pdf-g* with *Pdf-Gal4* resulted in complete loss of PDF from sLNvs but only partial loss from the lLNvs (Fig. 3A, B). The higher levels of PDF in lLNvs than in sLNvs is one possible explanation, but persistent PDF in lLNvs might also reflect the duration of expression of the CRISPR tools in the different cells. The sLNvs have an early larval origin, well before the lLNvs appear in the mid-late pupal stage (48). This temporal pattern is recapitulated by *Pdf-Gal4* (7), allowing sLNvs a much longer time for mutagenesis and subsequent protein turnover.

PDF is a key circadian neuropeptide in *Drosophila*, and the small PDF neurons (sLNvs) are critical for circadian behavior (14, 15). This is because PDF from the sLNvs is a key regulator of rhythmicity in constant darkness (13). The results here show that loss of PDF from these neurons also has a major effect on sleep structure, namely, a decrease in p-wake. Loss of PDF from the lLNvs has a qualitatively similar but much smaller quantitative decrease in p-wake (Fig. 3I-K). This indicates that PDF from both sources is wake-promoting, consistent with the fact that activation of lLNvs mediates arousal (18, 49). A greater role for lLNvs has been described in promoting wake through GABA signaling under light-dark conditions (68, 69). It is therefore possible that the lLNvs play a larger role in maintaining wakefulness under LD conditions, whereas rhythmic PDF release from the sLNvs in constant darkness promotes wake as well as maintains rhythmicity.

Two independent methods indicate that PDF functions adult-specifically to regulate rhythmicity and sleep. Although both worked effectively, the two strategies have distinguishing features. The inducible Cas9 system was very robust i.e., there was no leaky expression, which provides high confidence in the adult-specific conclusion. However, the induction was incomplete, leading to some mosaicism and considerable variability in the resulting phenotypes (Fig. S7A). Perhaps this could be solved by further altering the heat shock regimen, especially for rhythmicity as this assay is insensitive to the brief heat shock. Effects on sleep were more problematic, as even sleep in the control flies was impacted by heat shock (Fig. 4F). It is presently uncertain if the effect of heat shock alone on sleep structure is strain-specific or a more general effect.

In contrast to heat shock, the AGES system was highly efficient but exhibited some background mutagenesis in the absence of auxin at least with the PDF-Gal4 driver. As the AGES system was highly effective at inhibiting expression of a fluorescent protein marker (Fig. 5A, S8B), the background mutagenesis likely reflects the ability of even very low levels of Cas9 and gRNA expression to induce mutagenesis as well as the sensitivity of this assay. The lack of a similar leaky phenotype with the CLK856-Gal4 driver (Fig. 5G) suggests that it is Gal4-specific. Perhaps the CLK856-Gal4 driver only labels the PDF neurons at a later developmental stage, allowing less time to effect the leaky phenotype. Although mutagenesis sensitivity is a potential liability in this specific context, it is a generally positive feature of the CRISPR-based system: it does not require that drivers be highly expressed to mediate efficient mutagenesis. In the context of AGES, the use of lower expression Cas9 variants (33) might circumvent this issue. Importantly, auxin exposure alone had no detectable effect on circadian or sleep behavior (Fig. 5E, F). Thus, we find auxin exposure to be a less disruptive perturbation than heat shock and hence a mode of CRISPR induction more compatible for behavior analyses. Moreover, temporal regulation via AGES allows combining with other UAS transgenes for experiments like complementation analysis. Although the current version of AGES is incompatible with the many Gal80-insensitive split-Gal4 lines (25), it can be used with new Gal80-sensitive split-Gal4 lines (70). In summary, both temporal methods are effective, and we hope this exploration of their differences will help researchers choose the one best suited to their needs.

## Materials and Methods

### Fly Stocks and Rearing

All flies were raised on a standard cornmeal media supplemented with yeast in a temperature controlled incubator at 25°C in 12:12 Light:Dark cycles. The genotypes of all fly strains used in this study are listed in supplementary Table S3. Details on fly lines generated in this study are described in SI Appendix, Materials and Methods, Generation of Fly Lines.

### Locomotor Activity and Sleep Behavior

For behavior analysis of flies with throughout (not adult-specific) perturbations, 0-3 days old male flies of the appropriate genotype were collected and aged till they were about 2 weeks old at 25°C in 12:12 LD, with food changes every 3-4 days. The flies were then loaded into behavior tubes containing food (4% sucrose and 2% agar) and loaded onto *Drosophila* Activity Monitors (DAM; TriKinetics Inc., Waltham, MA). These DAMs were placed in light boxes with programmable LED light intensities inside a temperature-controlled incubator set to 25°C. Flies were entrained for 2-3 days with 12h:12h lights ON (500 lux): lights OFF before being switched to either constant darkness or constant light (500 lux). Circadian rhythmicity and sleep analysis was performed using the 2020 version of Sleep and Circadian Analysis MATLAB Program (SCAMP) developed by Christopher G. Vecsey (71).Details of locomotor behavior analysis are described in SI Appendix, Materials and Methods, Locomotor Activity and Sleep Behavior.

### Immunohistochemistry

Whole flies of the same genotype and age as the behavior experiment cohort were fixed in 4% paraformaldehyde. Dissected brains were washed and blocked with 10% Normal Goat Serum (NGS; Jackson labs). The following primary antibodies were used diluted in blocking buffer– chicken anti-GFP (Abcam ab13970; 1:1000), mouse anti-PDF (DSHB-PDF C7; 1:1000), guinea pig anti-VRI (gift of Paul Hardin; 1:3000 (72)), rat anti-TIM (Rosbash lab; 1:250), rabbit anti-dsRed (Takara Bio-632393; 1:300), rabbit anti-PER (Rosbash lab, 1:1000), rat anti-CRY (Rosbash lab, 1:200) and rat anti-RFP (Proteintech-5f8; 1:1000). Further details on image acquisition and analysis can be found in SI Appendix, Materials and Methods, Immunohistochemistry and Image Analysis.

See also, SI Appendix, Materials and Methods, Data representation and Statistics.

## Acknowledgements

We thank Michelle Lin for assistance with maintaining fly stocks. We thank Matthias Schlichting and all members of the Rosbash lab for useful discussions. We thank Paul Garrity, Justin Blau and Katharine Abruzzi for helpful comments on the manuscript. We thank Paul Hardin for the VRI antibody. We thank Heather Dionne, Aljoscha Nern and Gerry Rubin (Janelia Research campus) for the split-Gal4 lines, and Filip Port and Michael Boutros for the STOP-Cas9 flies. Stocks obtained from the Bloomington Drosophila Stock Center (NIH P40OD018537) and Vienna Drosophila Resource Center (VDRC) were used in this study. This work was supported by the Howard Hughes Medical Institute (HHMI).

## SI Materials and Methods

### Generation of Fly Lines

To generate lines expressing 3X-guides under UAS-control, we used previously described protocols (Port and Bullock, 2016; Schlichting et al., 2019). Briefly, gRNAs specific to the target genes were identified using the CRISPR Optimal Target Finder tool (Gratz et al., 2014). These gRNA sequences were incorporated into the tRNA backbone using 2 PCR reactions, primers for which are listed in supplementary Table S4. The PCR products were assembled into the UASpCFD6 vector (Addgene #73915) digested with BbsI-HF enzyme (NEB) using Gibson assembly. Sequence verified plasmids were injected into the attP2 site on the third chromosome by Rainbow Transgenic Flies Inc (Camarillo, CA, USA). Positive transformants were screened for by red eye color based on the miniwhite marker.

### Locomotor Activity and Sleep Behavior

Circadian rhythmicity and sleep analysis was performed using the 2020 version of Sleep and Circadian Analysis MATLAB Program (SCAMP) developed by Christopher G. Vecsey (Donelson et al., 2012). For period analysis, only rhythmic flies (Rhythmicity Index > 0.25) were used, apart from PDF perturbations in Fig. S6 and Table S2, where flies with Rhythmicity Index >0.2 were defined as rhythmic to look for residual short period rhythms in partially arrhythmic flies. E-peak timing was calculated as the highest point in bins from ZT6.5-ZT10.5 of average activity in 30 min bins exported from SCAMP. Sleep structure analysis for p-wake and p-doze (Wiggin et al., 2020) was also performed using this new version of SCAMP.

For phase shift experiments about 2-week-old males flies were entrained to LD for at least 3 days and on the last night of LD, 1 hour light pulse (500 lux) was delivered either at ZT15, ZT18 or ZT21. For one set of flies, no light pulse was given, and these were used as controls to calculate phase shift using the SCAMP program. Average shifts from days 3-5 are reported.

### Heat shock for inducible Cas9

For experiments with the inducible Cas9 system, 2-5 days old male flies of the appropriate genotype were heat shocked on 3 consecutive days. The first heat shock was 25 mins, and the two subsequent ones were 40 mins each. Heat shock was delivered by placing flies in an empty food vial and submerging the vial in a water bath set at 37°C. Flies were removed promptly and allowed to recover briefly at room temperature before being transferred to a fresh vial with standard cornmeal food. Once the flies recovered completely, the flies were placed in an incubator set to 25°C. Flies were aged for 2 weeks from the day of the first heat shock before loading them in behavior experiments. Control flies were collected at the same time and aged simultaneously but without the heat shock.

### Auxin feeding for adult-specific behavior with AGES

For AGES experiments, 2-5 days old male flies were fed Instant *Drosophila* Medium (Formula 424®, Carolina) reconstituted either in 10mM Auxin (GK2088, Glentham Life Sciences) or distilled water (for controls). Flies were fed this food for about 2 weeks before loading them in behavior tubes with standard behavior food (4% sucrose and 2% agar). Since CRISPR based mutagenesis irreversibly alters the DNA in the cells of interest, we used standard behavior food for behavior analysis post auxin feeding to make experiments simpler. For AGES time course, 0-2 day old male flies expressing gRNA and Cas9 with CLK856 Gal4 and AGES were collected and divided into 4 groups – No auxin, auxin fed for 2days, 4 days and 7 days. Flies were fed auxin for the indicated duration, starting later for shorter durations, and then tested for behavior on standard behavior food.

### Immunohistochemistry and Image Analysis

Whole flies of the same genotype and age as the behavior experiment cohort were fixed at ZT18-19 in 4% paraformaldehyde in PBS with 0.4% Triton X-100 for 2 hours and 30 minutes at room temperature. Tubes were wrapped in tin foil to protect them from light while rotating on a shaker. Flies were then washed three times (10 minutes each) with 0.4% PBST before dissecting the brain. Dissected brains were washed and blocked with 10% Normal Goat Serum (NGS; Jackson labs) in 0.4% PBST (blocking buffer) for 2 hours at room temperature or at 4°C overnight. The following primary antibodies were used diluted in blocking buffer– chicken anti-GFP (Abcam ab13970; 1:1000), mouse anti-PDF (DSHB-PDF C7; 1:1000), guinea pig anti-VRI (gift of Paul Hardin; 1:3000 (Glossop et al., 2003)), rat anti-TIM (Rosbash lab; 1:250, (Ma et al., 2021)), rabbit anti-PER (Rosbash lab, 1:1000, (Ma et al., 2021)), rabbit anti-dsRed (Takara Bio-632393; 1:300) and rat anti-RFP (Proteintech-5f8; 1:1000). Primary antibody was incubated for 24-36 hours at 4°C. The brains were then washed three times (30 minutes each) in 0.4% PBST, and corresponding secondary antibodies were used at a 1:500 dilution in blocking buffer and incubated at 4°C overnight. Brains were again washed three times (30 minutes each) in 0.4% PBS, mounted in VectaShield mounting medium (Vector Laboratories).

For CRY immunostaining, flies were kept in constant darkness for several days to allow accumulation and detection of CRY as described earlier (Yoshii et al., 2008). Flies were fixed at CT0-1 (DD4) in dark conditions. The rat anti-CRY antisera (Emery et al., 1998) were incubated with fixed heads of the null mutants, *cry^01^*(Dolezelova et al., 2007) to reduce background staining. This primary rat anti-CRY antibody was used at 1:200.

Images were acquired using Leica Stellaris 8 confocal microscope equipped with a white light laser. Images for one set of experiments were acquired using the Leica SP5 confocal microscope. For each experiment, the laser settings were kept constant across genotypes. Either 20X air objective or a 40X oil objective (for image quantification) with NA of 1.43 was used. Image analysis was performed with Fiji (Schindelin et al., 2012) using the Time Analyzer V3 plugin to mark ROIs across channels and measure intensity of the PDF channel. Background intensity for a z-stack was measured from three random background regions, averaged, and subtracted from each ROI.

### Data representation and Statistical Analysis

Some of the figure panels (Fig. S5A, S7A and S8A) were created using BioRender. Boxplots were generated using BoxPlotR (Spitzer et al., 2014). For all boxplots, center lines show the medians; box limits indicate the 25th and 75th percentiles as determined by R software; whiskers extend 1.5 times the interquartile range from the 25th and 75th percentiles, outliers are represented by dots. Source data for all figures are included in SI Dataset 1. Statistical tests performed are described in the figure legends. All statistical tests were performed using the free online tool Statistics Kingdom https://www.statskingdom.com/index.html

### Supplementary Figures

**Fig. S1:**
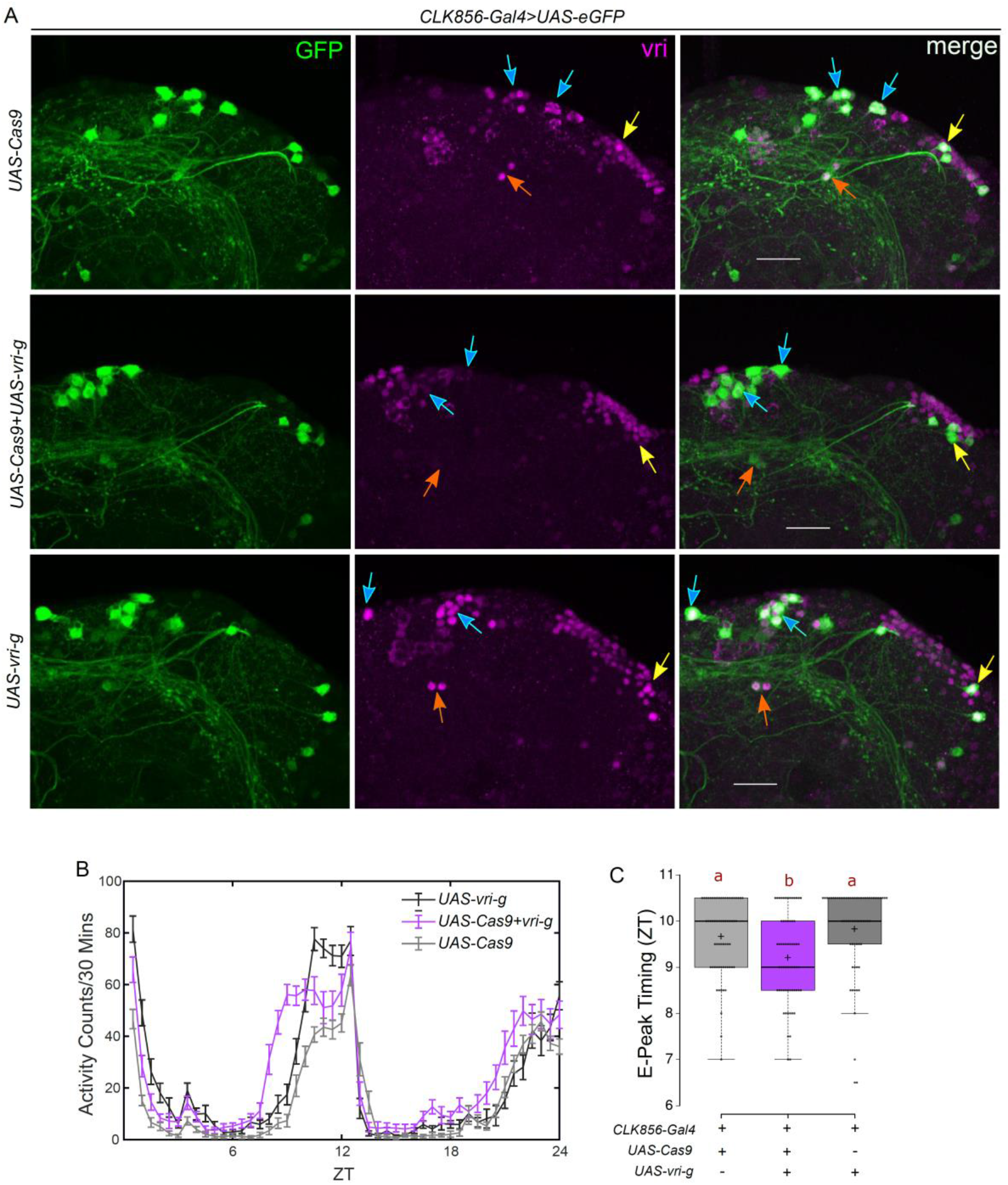
Loss of VRI from clock neurons leads to an advanced evening peak. **A**. Representative images of the dorsal circadian neurons stained for GFP and VRI at ZT19. There is loss of VRI staining in cells expressing *CLK856-Gal4* (marked by GFP) but no loss in GFP-negative cells. Arrows point to examples of Gal4-positive DN1s (blue), DN2s (red) and DN3s (yellow) in the indicated genotypes. Scale bar represents 20µm. **B**. Average activity counts for flies in an experiment plotted as a function of time. Flies with loss of VRI in *CLK856-Gal4* labelled neurons showed an advanced evening peak. **C**. Quantification of the evening peak (E-peak) timing from individual flies, n ≥ 62 per genotype from at least two independent experiments. Letters represent statistically distinct groups; p<0.01, Kruskal Wallis test followed by a post hoc Dunn’s test.

**Fig. S2:**
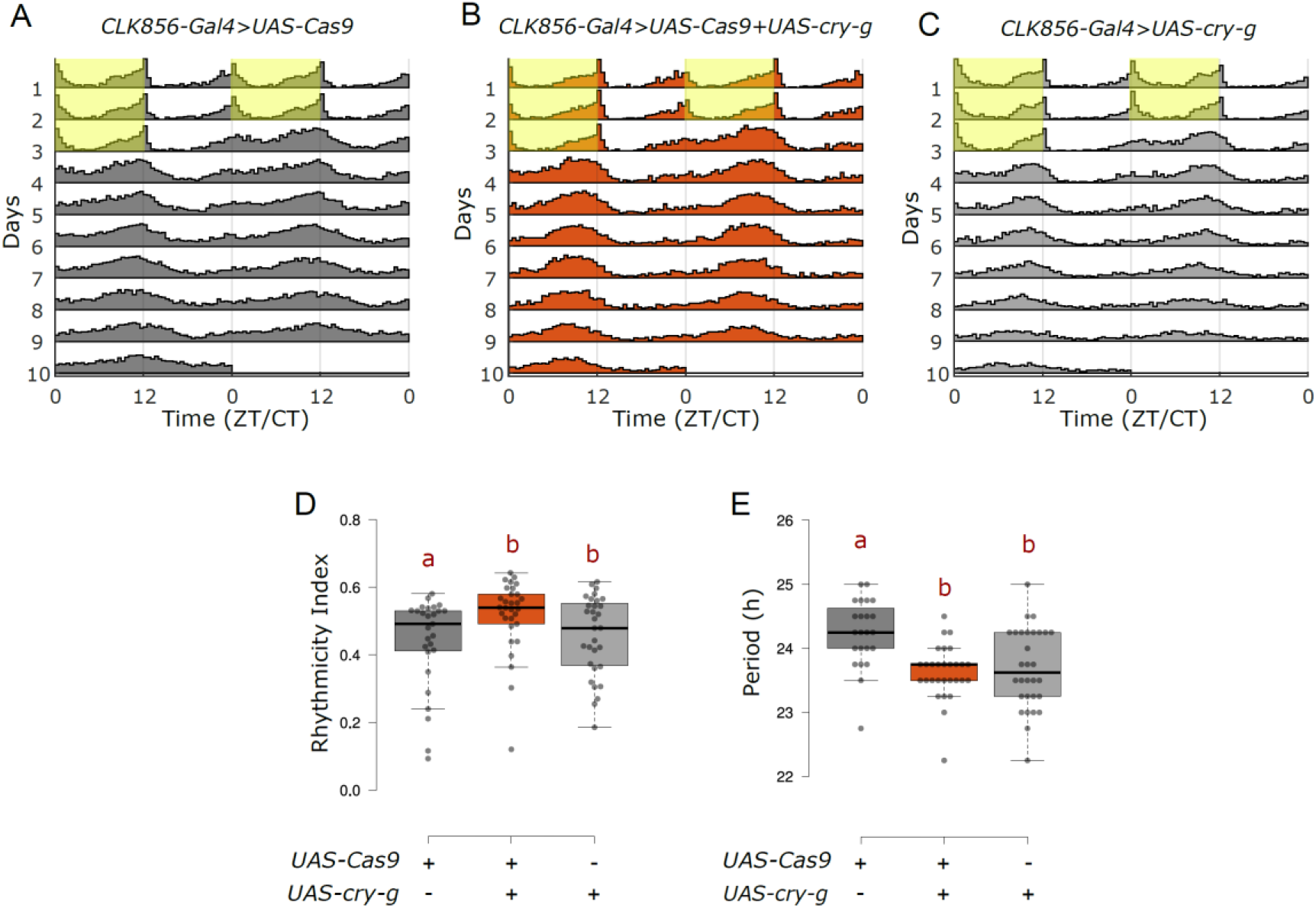
CRY mutation in clock neurons does not affect circadian behavior under constant darkness. **A-C**. Actograms represent double-plotted average activity of flies from an experiment across multiple days. Yellow panels indicate lights ON. **D**. Rhythmicity Index for individual flies quantified for DD2-7 represented by a boxplot, n ≥ 27 per genotype, letters represent statistically distinct groups, p<0.01, Mann-Whitney U test post Kruskal Wallis ANOVA. **E**. Free running period under constant darkness for individual flies quantified for DD2-7 represented by a boxplot, letters represent statistically distinct groups; p<0.01, Kruskal Wallis test followed by a post hoc Dunn’s test.

**Fig. S3:**
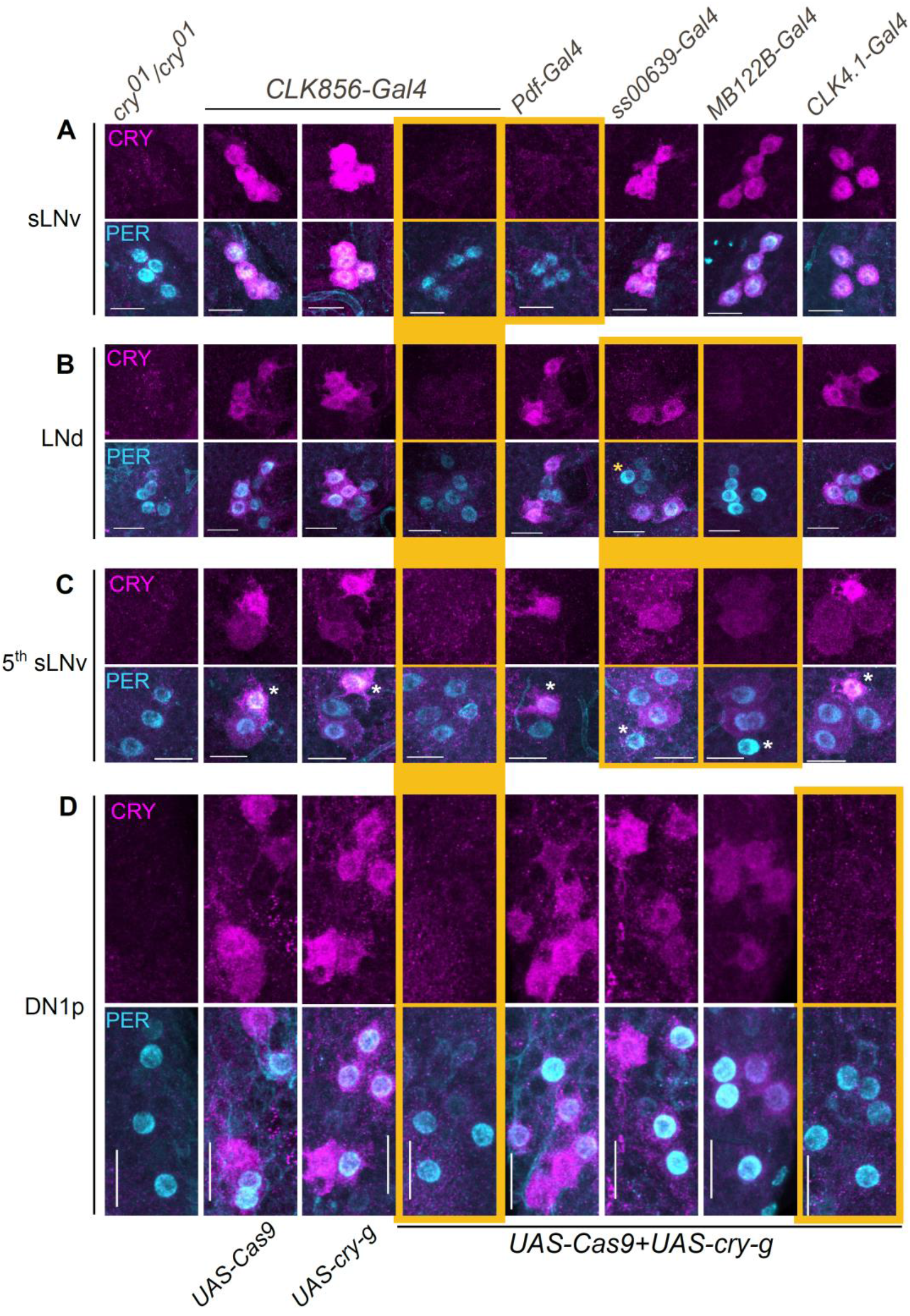
CRISPR mediated cell specific mutagenesis of cry leads to cell-type specific loss of CRY-staining. Images from brains co-stained with CRY and PER, to label clock neurons at CT0-1 (DD4). In controls (columns 2 and 3), CRY staining is detected strongly in the sLNvs, 3 of the 6 LNds, the 5^th^ sLNv and a subset of the DN1ps whereas weakly in the lLNvs. CRY staining was absent from all clock neurons in the cry null mutant (*cry^01^/cry^01^*; column 1). Similarly, CRISPR mediated knockout of CRY with *CLK856-Gal4* led to severely reduced CRY staining in all clock neurons (A-D; column 4), whereas CRISPR mutagenesis with specific Gal4s led to loss of CRY staining from specific neurons as described below. Panels where CRY staining is perturbed are highlighted in yellow. **A**. CRY staining in the sLNv neurons was perturbed only with *Pdf-Gal4.* **B**. CRY mutagenesis with *MB122B-Gal4* led to loss of CRY staining from all three LNds whereas that with *ss00639-Gal4* led to loss of CRY staining from only one LNd (marked by asterisk). **C.** In the 5^th^ sLNv (marked by asterisk), CRY staining is lost with CRISPR mutagenesis in *MB122B-Gal4* and *ss00639-Gal4.* **D**. CRY staining in the DN1ps is lost only when expression is targeted with the *CLK4.1-Gal4.* Scale bars represent 10µm.

**Fig. S4:**
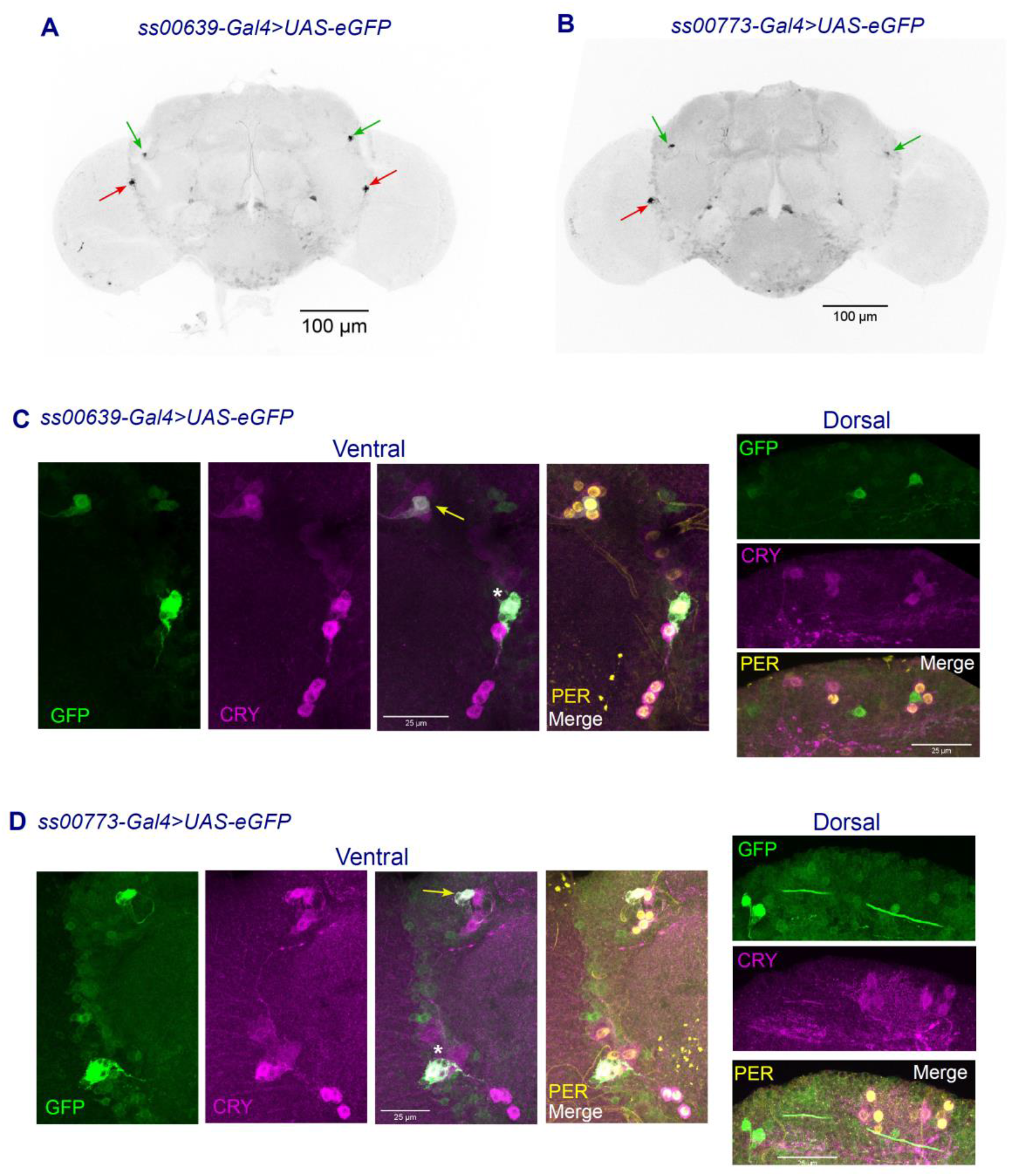
Split-Gal4 drivers labelling a subset of evening cells. **A-B:** Full brain expression patterns of the two split-Gal4 drivers labelled by UAS-eGFP. **C-D:** Split-Gal4 lines co-stained with CRY and PERIOD (PER) to label clock neurons. Both split-Gal4s label the CRY and PER positive 5^th^ sLNv (marked by asterisk) and one LNd (yellow arrow). In panels A-B, red arrows point to the 5^th^-sLNv and green arrows to an LNd neuron as identified by co-staining with clock proteins in C-D. The *ss00639-Gal4* labels the 5^th^-sLNv + 1 LNd per hemisphere in all brains. The *ss00773-Gal4* also labels the 5^th^-sLNv + 1 LNd, but it does so more variably and labels about 2-4 of these clock cells per brain. Both Gal4s also label a few extra cells, *ss00773-Gal4* more so than *ss00639-Gal4,* but none of them are either PER or CRY positive.

**Fig. S5:**
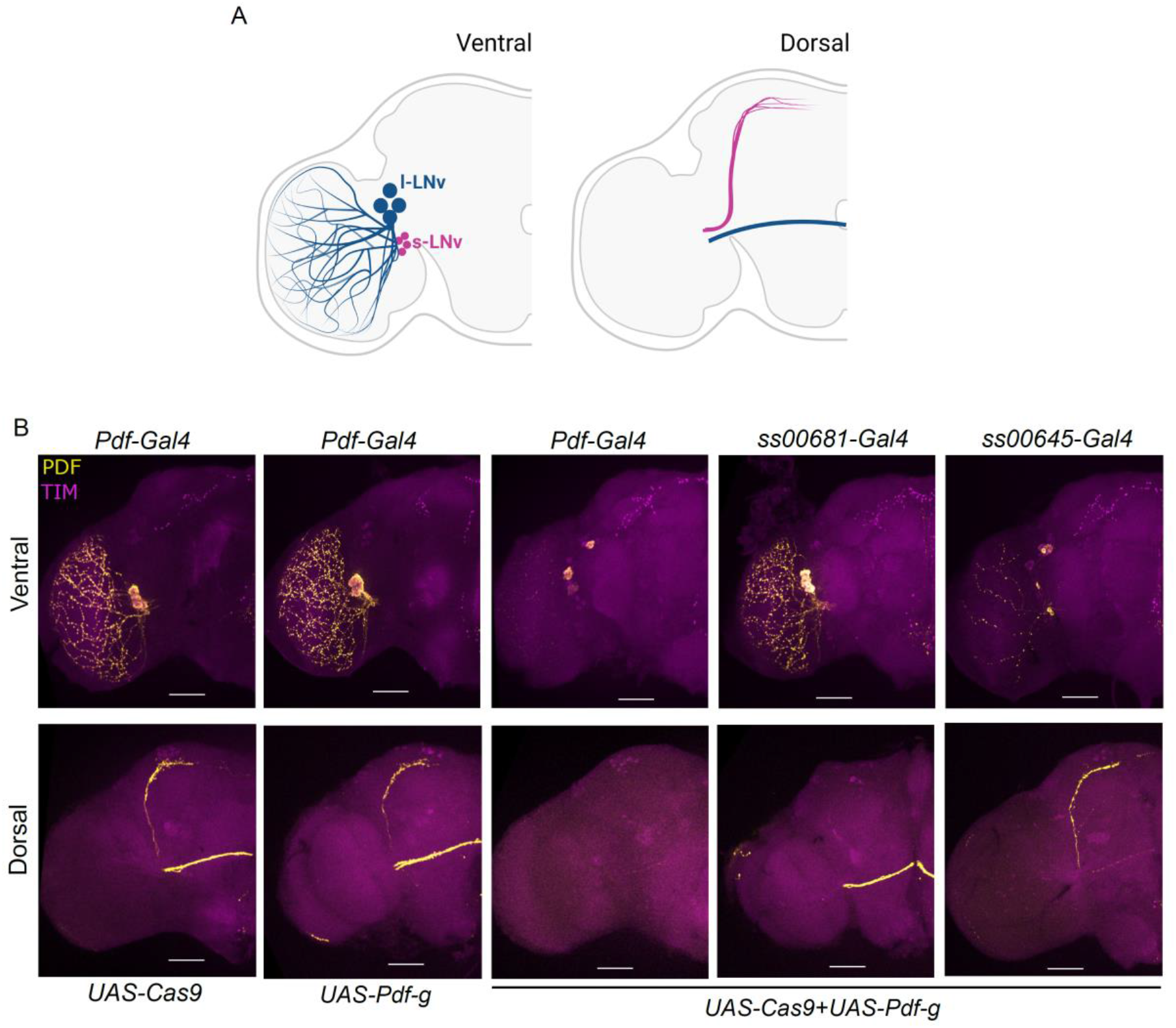
CRISPR strategy successfully eliminates PDF staining from cell type-specific projections. **A**. Cartoon representation of the projection patterns of small (magenta) and large (blue) LNV neurons in the ventral and dorsal *Drosophila* brain. **B**. Representative images of hemibrains imaged from the ventral (top) or dorsal (bottom) side stained for PDF and TIM. For the dorsal panels, brightness-contrast was enhanced for the PDF channel equally across genotypes to confirm absence of PDF staining from the respective processes. Expression of both *UAS-Cas9* and *UAS-Pdf-g* with *Pdf-Gal4* leads to loss of all PDF staining from all PDF neurons. Expression with *ss00681-Gal4* that labels only the sLNvs leads to loss of PDF staining from their cell bodies and projections with no obvious effect on PDF staining in lLNvs. Expression with *ss00645-Gal4* leads to loss of PDF staining in most projections of these neurons with no obvious effect on PDF staining in sLNvs.

**Fig. S6:**
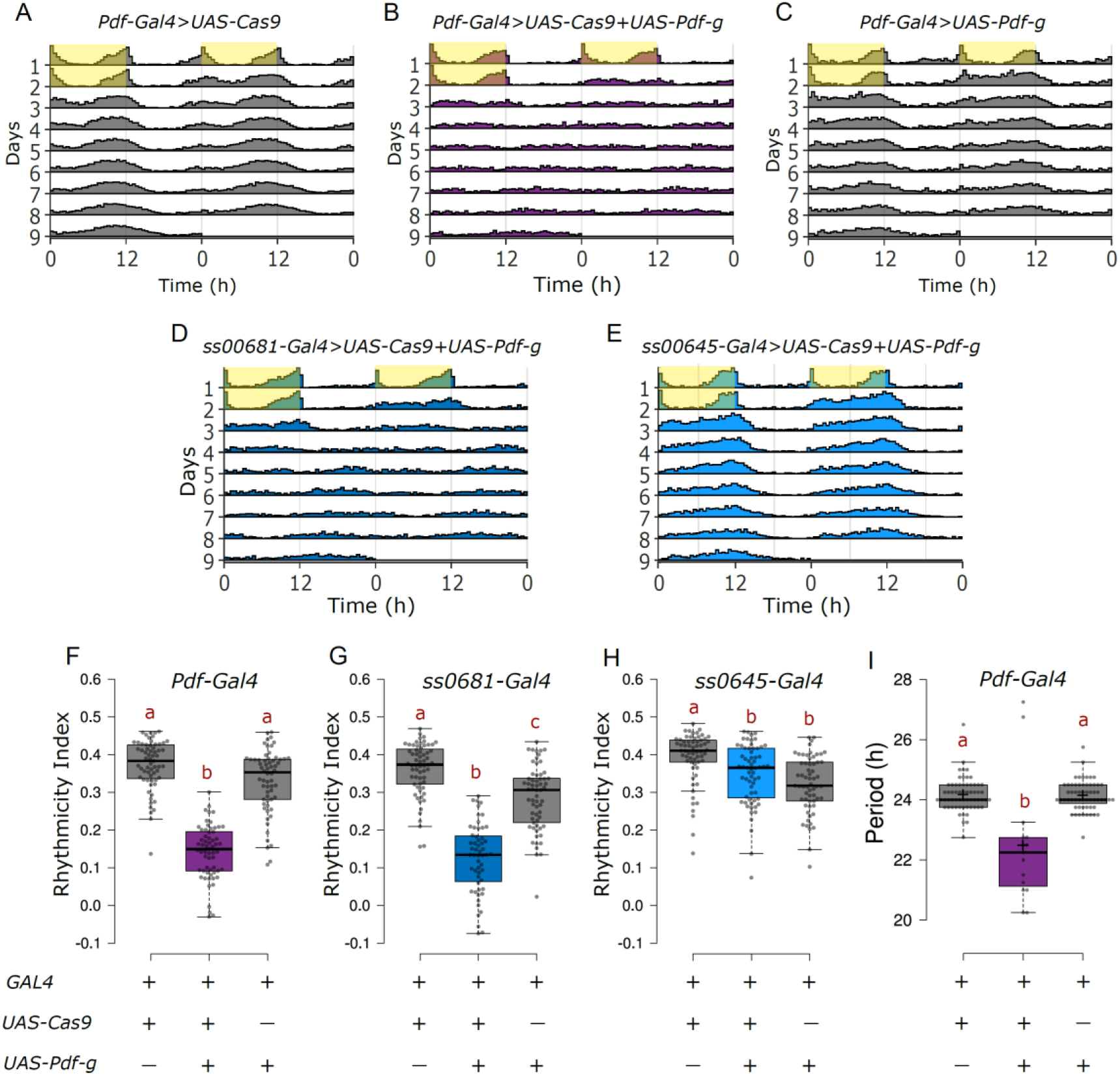
Loss of PDF from sLNvs leads to loss of rhythmicity under constant darkness. **A-E**. Actograms represent double-plotted average activity of flies from an experiment across multiple days. Yellow panels indicate lights ON. **F-H**. Rhythmicity Index for individual flies quantified for DD1-4 represented by a boxplot, n ≥ 55 per genotype from at least two independent experiments, letters represent statistically distinct groups; p<0.01, Kruskal Wallis test followed by a post hoc Dunn’s test. Loss of PDF from both sLNvs and lLNvs or from sLNvs alone causes arrhythmicity in constant darkness, whereas loss of PDF from lLNvs alone has no effect. **I.** Free running period for flies of the indicated genotypes, rhythmic flies (RI>0.2) with loss of PDF in PDF neurons have a shorter period.

**Fig. S7:**
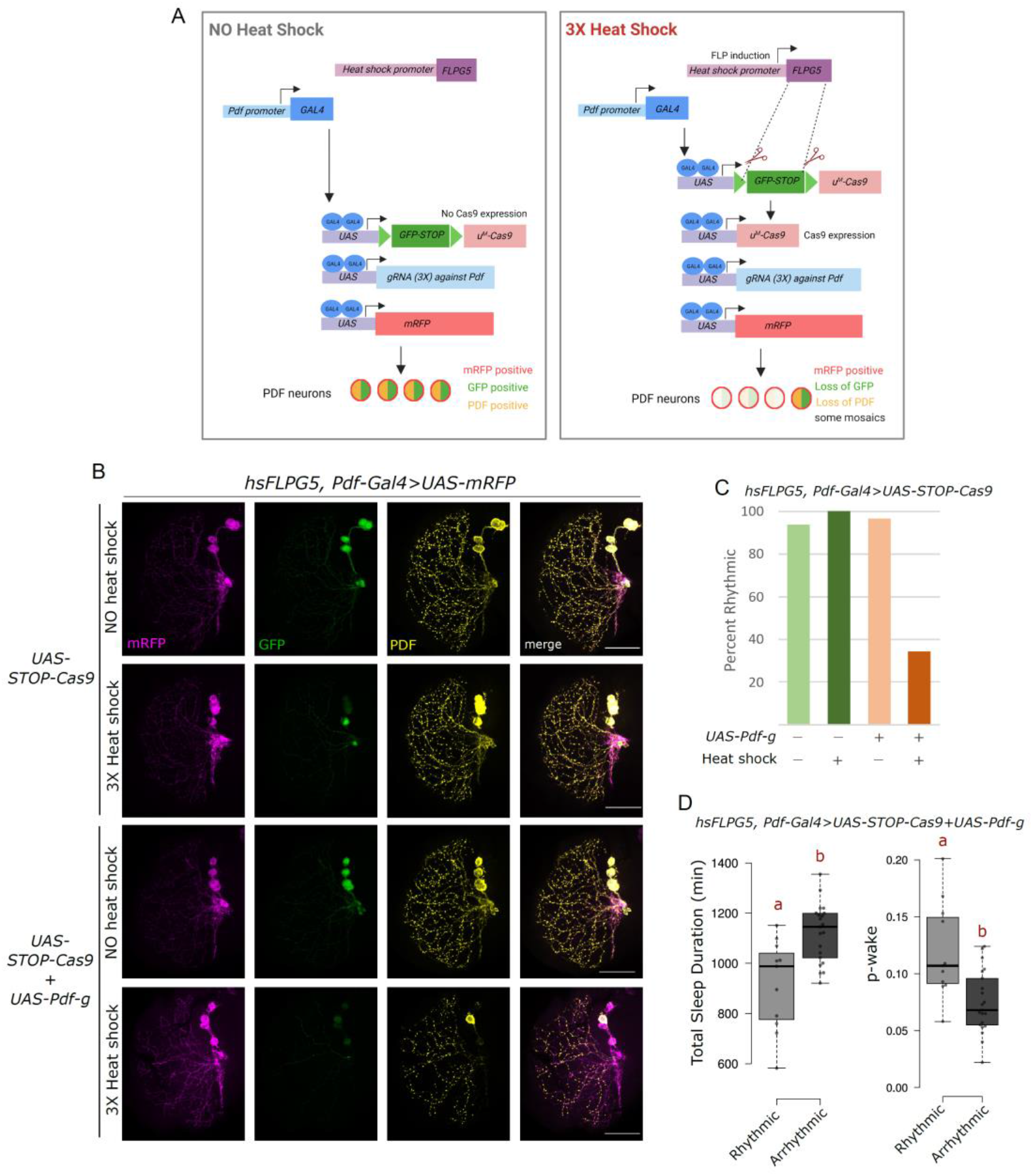
The inducible-Cas9 systems is an effective tool for adult specific and cell type specific CRISPR mutagenesis despite some mosaics. **A**. Cartoon representation of the inducible Cas9 system (Port et. al., 2020) adapted to the adult-specific perturbation of PDF from PDF neurons. **B**. Representative images of the optic lobe ventral projections of the lLNvs. Scale bar represents 50µm. **C**. Plot indicating the percentage of flies rhythmic under the indicated conditions. **D**. Sleep parameters of flies expressing both *UAS-Cas9* and *UAS-Pdf-g* with heat-shock separated into rhythmic (RI>0.25) and arrhythmic (RI<0.25) categories, n=11 for rhythmic and 21 for arrhythmic. Arrhythmic flies have more sleep and a lower p-wake; p<0.01, Mann Whitney U test. These rhythmic flies could be the result of the mosaicism, i.e., PDF expression in 1-2 sLNvs might be sufficient for quasi-normal circadian and sleep behavior.

**Fig. S8:**
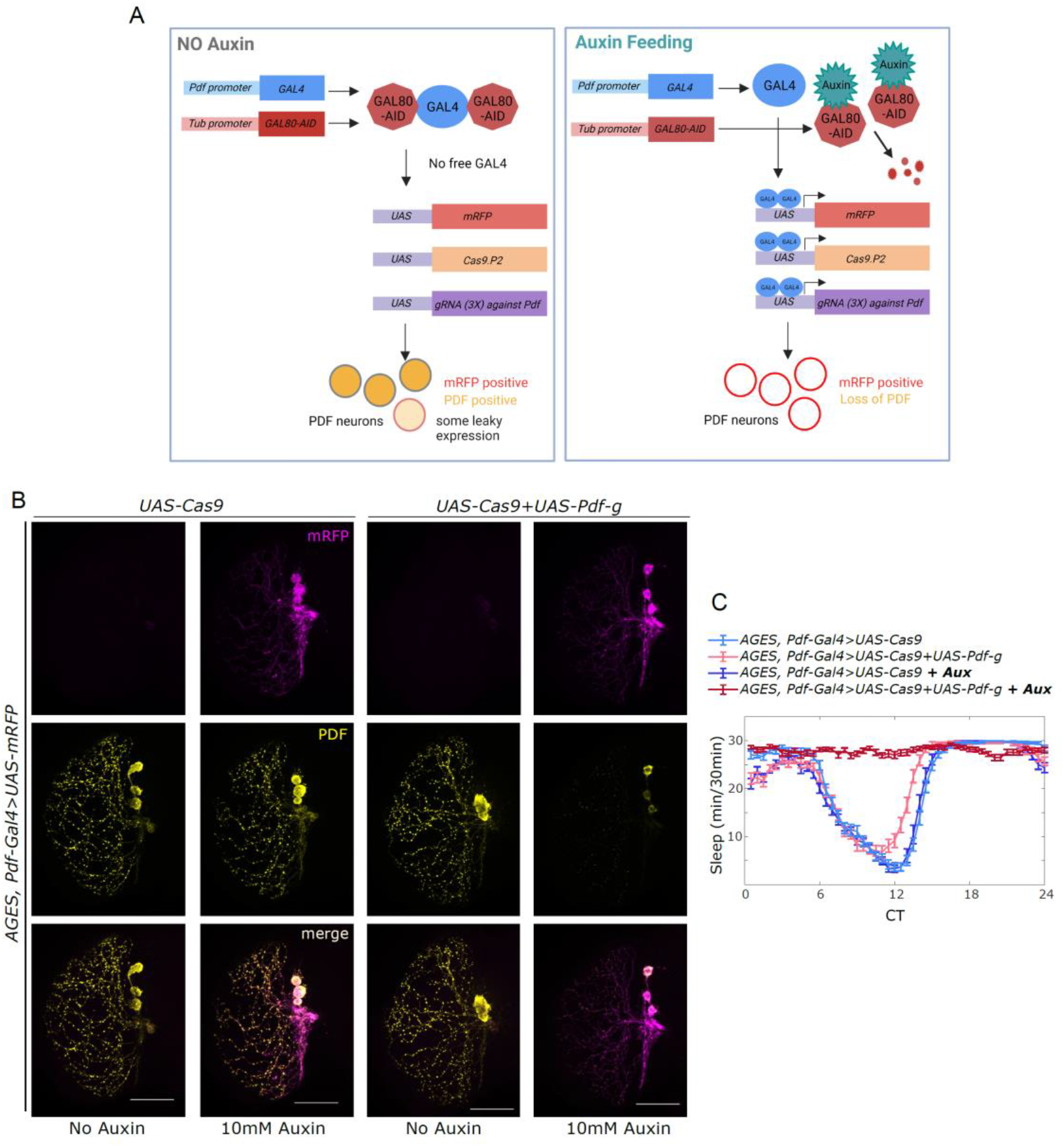
AGES is mildly leaky, but still an effective tool for adult-specific, cell type-specific CRISPR mutagenesis. **A**. Cartoon representation of the AGES system (McClure et. al., 2022) adapted to the adult specific perturbation of PDF from PDF neurons. **B**. Representative images of the optic lobe ventral projections of the lLNvs. Scale bar represents 50µm. **C**. Sleep plot representing average sleep of flies from DD1-4 in 30-minute bins.

### Supplementary Tables

**Table S1:** Summary of circadian behavior with cell type-specific CRY mutants in constant light.

| Genotype | Cells labelled | n | Rhythmicity Index LL2-7 (Mean ± SEM) | %Rhythmic (RI>0.25) | Period (h) (Mean ± SEM) |
| --- | --- | --- | --- | --- | --- |
| *CLK856-Gal4>UAS-Cas9* | Most clock neurons (many DN3s missing) | 85 | 0.13 ± 0.01 | 10.6 | - |
| *CLK856-Gal4>UAS-CRYg* | | 72 | 0.11 ± 0.01 | 6.9 | - |
| ***CLK856-Gal4>UAS-Cas9+UAS-CRYg*** |  | **92** | **0.40 ± 0.01** | **90.2** | **24.71 ± 0.07** |
| *PDF-Gal4>UAS-Cas9* | sLNvs+ ILNvs | 118 | 0.10 ± 0.01 | 6.8 | - |
| *PDF-Gal4>UAS-CRYg* | | 119 | 0.12 ± 0.01 | 5.0 | - |
| ***PDF-Gal4>UAS-Cas9+UAS-CRYg*** |  | **121** | **0.17 ± 0.01** | **14.9** | - |
| *dvPDF-Gal4>UAS-Cas9* | sLNvs+ ILNvs+ 5<sup>th</sup> sLNv+ 1 CRY<sup>+</sup> ITP<sup>+</sup> LNd + 3 CRY-LNds | 56 | 0.14 ± 0.01 | 8.9 | - |
| *dvPDF-Gal4>UAS-CRYg* | | 51 | 0.15 ± 0.01 | 9.8 | - |
| ***dvPDF-Gal4>UAS-Cas9+UAS-CRYg*** |  | **58** | **0.52 ± 0.02** | **91.4** | **24.24 ± 0.06** |
| *MB122B-Gal4>UAS-Cas9* | 5<sup>th</sup> sLNv + 1 CRY<sup>+</sup>ITP<sup>+</sup> + 2 CRY<sup>+</sup> Trissin<sup>+</sup> LNds | 64 | 0.15 ± 0.01 | 9.4 | - |
| *MB122B-Gal4>UAS-CRYg* | | 61 | 0.14 ± 0.01 | 3.3 | - |
| ***MB122B-Gal4>UAS-Cas9+UAS-CRYg*** |  | **63** | **0.37 ± 0.02** | **79.4** | **25.08 ± 0.08** |
| *ss00639-Gal4>UAS-Cas9* | 5<sup>th</sup> sLNv + 1 CRY<sup>+</sup> LNd | 64 | 0.14 ± 0.01 | 9.4 | - |
| *ss00639-Gal4>UAS-CRYg* | | 58 | 0.15 ± 0.01 | 10.3 | - |
| ***ss00639-Gal4>UAS-Cas9+UAS-CRYg*** |  | **60** | **0.41 ± 0.01** | **91.7** | **25.11 ± 0.06** |
| *ss00773-Gal4>UAS-Cas9* | 5<sup>th</sup> sLNv + 1 CRY<sup>+</sup> LNd | 63 | 0.14 ± 0.01 | 4.8 | - |
| *ss00773-Gal4>UAS-CRYg* | | 60 | 0.15 ± 0.01 | 8.3 | - |
| ***ss00773-Gal4>UAS-Cas9+UAS-CRYg*** |  | **61** | **0.29 ± 0.02** | **60.7** | **25.06 ± 0.14** |
| *CLK4.1-Gal4>UAS-Cas9* | DN1p | 72 | 0.12 ± 0.01 | 9.7 | - |
| *CLK4.1-Gal4>UAS-CRYg* | | 69 | 0.13 ± 0.01 | 5.8 | - |
| ***CLK4.1-Gal4>UAS-Cas9+UAS-CRYg*** |  | **83** | **0.33 ± 0.01** | **74.7** | **21.82 ± 0.08** |

**Table S2:** Summary of circadian and sleep behavior upon loss of PDF from specific neurons.

| Genotype | cells labelled | n | Percent Rhythmic (RI>0.2) | Rhythmicity Index (DD1-4) | Average total sleep per day(min) | Activity while awake | pwake | pdoze |
| --- | --- | --- | --- | --- | --- | --- | --- | --- |
|  |  |  |  | Mean ± SEM |  |  |  |  |
| *PDF-Gal4>UAS-Cas9* | All sLNvs + ILNvs | 63 | 98.4 | 0.37 ± 0.01 | 914 ± 22 | 1.62 ± 0.04 | 0.12 ± 0.006 | 0.37 ± 0.01 |
| *PDF-Gal4>UAS-PDFg* |  | 62 | 90.3 | 0.33 ± 0.01 | 873 ± 23 | 1.79 ± 0.05 | 0.13 ± 0.007 | 0.35 ± 0.01 |
| ***PDF-Gal4>UAS-Cas9 + UAS-PDFg*** |  | **61** | **24.6** | **0.14 ± 0.01** | **1178 ± 19** | **2.52 ± 0.10** | **0.06 ± 0.005** | **0.40 ± 0.01** |
| *ss00681-Gal4>UAS-Cas9* | All sLNvs | 63 | 96.8 | 0.36 ± 0.01 | 924 ± 19 | 1.81 ± 0.04 | 0.12 ± 0.006 | 0.38 ± 0.01 |
| *ss00681-Gal4>UAS-PDFg* |  | 62 | 83.9 | 0.29 ± 0.01 | 972 ± 21 | 2.17 ± 0.07 | 0.12 ± 0.006 | 0.42 ± 0.01 |
| ***ss00681-Gal4>UAS-Cas9 + UAS-PDFg*** |  | **55** | **23.6** | **0.13 ± 0.01** | **1284 ± 13** | **2.57 ± 0.10** | **0.04 ± 0.003** | **0.44 ± 0.01** |
| *ss00645-Gal4>UAS-Cas9* | Variably 3-4 ILNvs per hemi-brain | 62 | 96.8 | 0.39 ± 0.01 | 957 ± 23 | 1.79 ± 0.05 | 0.11 ± 0.008 | 0.34 ± 0.01 |
| *ss00645-Gal4>UAS-PDFg* |  | 61 | 96.7 | 0.32 ± 0.01 | 1068 ± 14 | 2.37 ± 0.10 | 0.08 ± 0.004 | 0.38 ± 0.01 |
| ***ss00645-Gal4>UAS-Cas9 + UAS-PDFg*** |  | **63** | **95.2** | **0.35 ± 0.01** | **1142 ± 15** | **2.20 ± 0.08** | **0.07 ± 0.003** | **0.38 ± 0.1** |

**Table S3:** List of fly strains.

| **Genotype** | **Reference/Source** | **Stock number** |
| --- | --- | --- |
| CLK856-Gal4 | (Gummadova et al., 2009);<br>Bloomington Drosophila Stock Center | BDSC 93198 |
| UAS-eGFP | Bloomington Drosophila Stock Center | BDSC 5430 |
| UAS-mRFP | Bloomington Drosophila Stock Center | BDSC 32219 |
| UAS-Cas9.P2 (attP40) | Bloomington Drosophila Stock Center | BDSC 58985 |
| UAS-Cas9.P2 (attP2) | Bloomington Drosophila Stock Center | BDSC 58986 |
| AGES | (McClure et al., 2022);<br>Bloomington Drosophila Stock Center | BDSC 92470 |
| hsFLPG5 | Bloomington Drosophila Stock Center | BDSC 55814 |
| *cry*<sup>01</sup> | (Dolezelova et al., 2007) | NA |
| PDF-Gal4 | (Renn et al., 1999) | NA |
| CLK4.1-Gal4 | (L. Zhang et al., 2010; Y. Zhang et al., 2010) | NA |
| dvPDF-Gal4 | (Bahn et al., 2009) | NA |
| MB122B-Gal4 | (Bulthuis et al., 2019) | Lines made by<br>Drs. Dione, Nern<br>and Rubin, Janelia<br>Research Campus |
| ss00681-Gal4 | (Liang et al., 2017) |  |
| ss00645-Gal4 | (Schlichting et al., 2019) |  |
| ss00639-Gal4 | NA |  |
| ss00773-Gal4 | NA |  |
| UAS-STOP-Cas9 (UAS-FRT-mEGFP-FRT-uM-Cas9) | (Port et al., 2020);<br>Vienna Drosophila Resource Center | VDRC 340008 |
| UAS-*per-g* | (Schlichting et al., 2019) | NA |
| UAS-3X-vrig (attP2) | This study | NA |
| UAS-3X-cryg (attP2) | This study | NA |
| UAS-3X-Pdfg (attP2) | This study | NA |

**Table S4:**
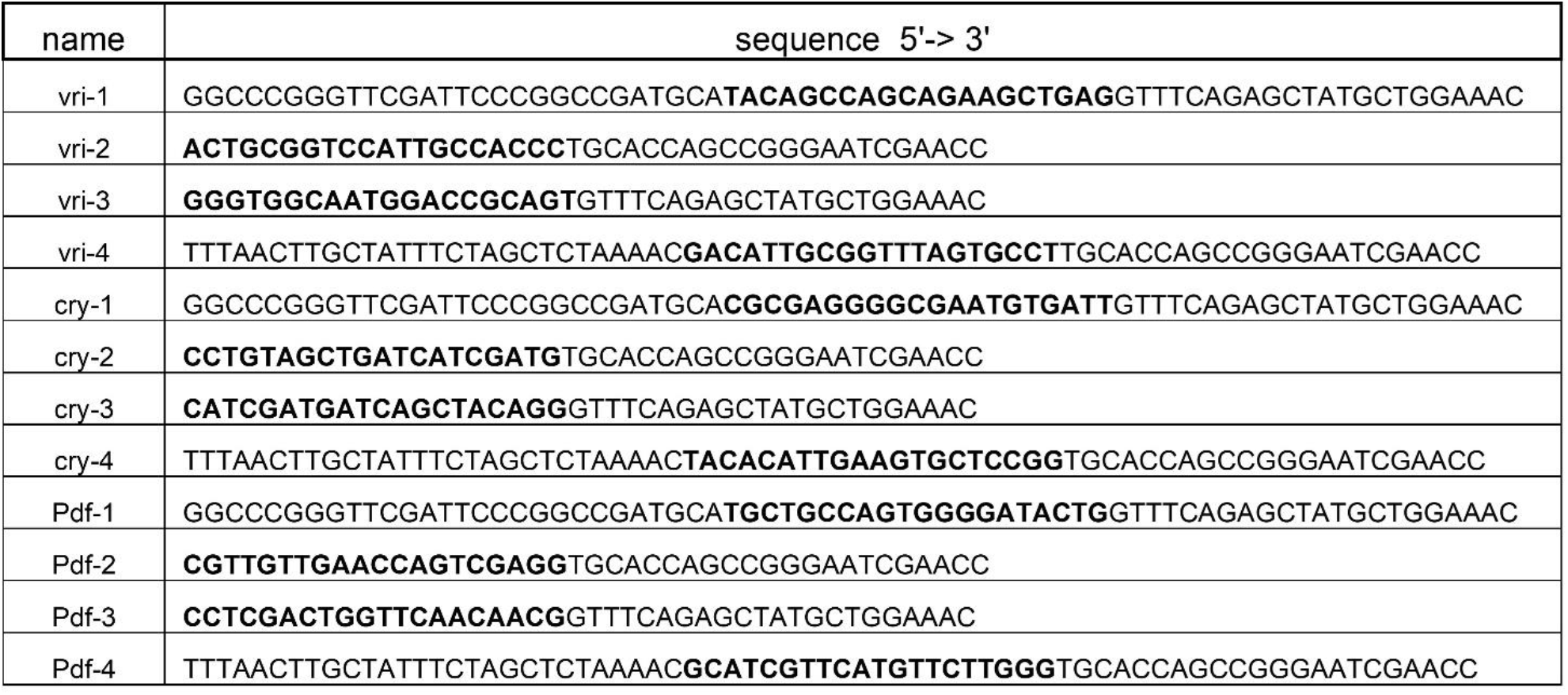
Primer sequences used for cloning.

**Legend for Supplementary Dataset 1:** File contains source data and statistical test for all figures.

